# Estimating effective connectivity in neural networks: comparison of derivative-based and correlation-based methods

**DOI:** 10.1101/2024.02.05.578871

**Authors:** Niklas Laasch, Wilhelm Braun, Lisa Knoff, Jan Bielecki, Claus C. Hilgetag

## Abstract

Inferring and understanding the underlying connectivity structure of a system solely from the observed activity of its constituent components is a challenge in many areas of science. In neuroscience, techniques for estimating connectivity are paramount when attempting to understand the network structure of neural systems from their recorded activity patterns. To date, no universally accepted method exists for the inference of effective connectivity, which describes how the activity of a neural node mechanistically affects the activity of other nodes. Here, focussing on purely excitatory networks of small to intermediate size and continuous node dynamics, we provide a systematic comparison of different approaches for estimating effective connectivity. Starting with the Hopf neuron model in conjunction with known ground truth structural connectivity, we reconstruct the system’s connectivity matrix using a variety of algorithms. We show that, in sparse non-linear networks with delays, combining a lagged-cross-correlation (LCC) approach with a recently published derivative-based covariance analysis method provides the most reliable estimation of the known ground truth connectivity matrix. We outline how the parameters of the Hopf model, including those controlling the bifurcation, noise, and delay distribution, affect this result. We also show that in linear networks, LCC has comparable performance to a method based on transfer entropy, at a drastically lower computational cost. We highlight that LCC works best for small sparse networks, and show how performance decreases in larger and less sparse networks. Applying the method to linear dynamics without time delays, we find that it does not outperform derivative-based methods. We comment on this finding in light of recent theoretical results for such systems. Employing the Hopf model, we then use the estimated structural connectivity matrix as the basis for a forward simulation of the system dynamics, in order to recreate the observed node activity patterns. We show that, under certain conditions, the best method, LCC, results in higher trace-to-trace correlations than derivative-based methods for sparse noise-driven systems. Finally, we apply the LCC method to empirical biological data. Choosing a suitable threshold for binarization, we reconstruct the structural connectivity of a subset of the nervous system of the nematode *C. Elegans*. We show that the computationally simple LCC method performs better than another recently published, computationally more expensive reservoir computing-based method. We apply different methods to this dataset and find that they all lead to similar performances. Our results show that a comparatively simple method can be used to reliably estimate directed effective connectivity in sparse neural systems in the presence of spatio-temporal delays and noise. We provide concrete suggestions for the estimation of effective connectivity in a scenario common in biological research, where only neuronal activity of a small set of neurons, but not connectivity or single-neuron and synapse dynamics, are known.

## 1 Introduction

### 1.1 Connectivity of neural circuits

What can be learnt about the network structure of a neural system from observations of the dynamics of its constituent components? In network neuroscience, three principal modes of connectivity can be distinguished [1], [2], specifically structural, functional, and effective connectivity (SC, FC, and EC, respectively). The main aim of this paper is to compare approaches to estimate EC, which involves SC as a first step. Hence, we first explore and compare approaches that aim to estimate the SC, and finally, with knowledge of the nodal dynamics, the EC of a network-based neural system. Therefore, for the purposes of this paper, we define EC to consist of connectivity information in the form of SC plus a model for the nodal dynamics. Without known ground-truth (GT) SC, a given EC can be validated by comparing the simulated activity time series with recorded ones. This validation step constitutes another important aspect of this paper.

To start, SC [3] refers to the actual measured anatomical connectivity among neural elements, for example by white matter tracts. In humans, diffusion tensor imaging is used to approximate SC [4]. Obtaining reliable and detailed information on SC requires invasive experiments, which are prohibitive in humans [5]. In fact, the only adult organism with completely known SC is the nematode *C*.*elegans* [6], [7]. Recently, also the SC of the larval stage of the fruit fly *Drosophila* has become available [8]. We state from the outset that SC can be given by weighted or binary connectivity matrices from a mathematical point of view, resulting in connectomes that can, in general, be represented by weighted directed graphs. In particular, binary (or unweighted) connectivity matrices can be obtained from weighted ones by application of a suitable threshold.

Second, FC [9], [10] is the most commonly considered connectivity mode, for example in the context of resting-state functional magnetic resonance imaging (fMRI), but also for electroencephalography (EEG) and magnetoencephalography (MEG) recordings. By using correlation measures, this mode comprises statistical dependencies between brain regions that are not necessarily spatially close or structurally connected to each other. Because no dynamical model for the nodes is needed to compute FC, this method is model-free: only node time series are required to compute FC. Finally, EC [11], [12] estimates causal connectivity, via which the activity in one neural node mechanistically affects the activity in other nodes. Standard methods to obtain EC are Granger causality, dynamic causal modeling, Bayesian networks and transfer entropy [1]. In addition, there are also perturbation-based methods to infer EC, see [13], [14], [15]. In each case, EC is a model-based method that needs, in addition to SC, assumptions about the dynamics of each individual node. For linear systems, there are systematic ways to estimate EC, possibly constrained by SC, by optimization procedures [16], [17]. As such, in contrast to FC, it can explain the so-called network effect [17]: two nodes can be unconnected or weakly connected, but still be strongly correlated. There are many possibilities for this to occur, which we review in the following paragraphs, together with methods to estimate different types of connectivity.

### 1.2 Estimating different types of connectivity

A straightforward way to compute FC is to assign high connectivity to nodes whose time series co-fluctuate, hence showing a high (positive or negative) correlation, and assigning low connectivity to nodes that only have small correlation, possibly followed by thresholding. Correlations are a standard choice for FC when dealing with fMRI data with comparably low temporal resolution, but for temporally ‘richer’ signals like EEG/MEG, which exhibit a mixture of rhythms, filtering in defined spectral bands followed by computation of correlations is a more standard approach [18], [19]. Following this approach, one can compute a FC matrix [20], [21] and compare it to reconstructed anatomical connections (i.e., SC) forming the structural connectome [5]. The FC approach, although it is traditionally used in network neuroscience due to its mathematical simplicity, has a number of shortcomings. Because correlation (normalized zero-lag covariance) is intrinsically symmetric, estimates of FC based on pairwise correlations result in symmetric connectivity matrices. Moreover, activity correlations may not necessarily reflect structural connectivity [22]. In particular, FC does not necessarily reflect static SC, e.g. [23], [24]. One main reason for this discrepancy is that EC or FC can be estimated for different behavioral states (e.g. when the subject is at rest compared to when it performs a certain task), whereas SC is unique to each subject, and changes on a much slower timescale, if at all. A consequence is that there may be states during which a physically present connection is down-modulated, and hence, an SC estimate recorded using that state would result in a missing connection. One example is given by EC modulation during passive movie viewing inferred from fMRI data [22], which was shown to be mainly due to changes in the local activity caused by the increased stimulus load, and not by SC changes. In another scenario, two structurally uncoupled nodes show high correlation either because of common input from a third shared neighbor node (indirect coupling), or from outside of the network (common input). This scenario is one example of a confounder motif [22]. Because high correlation is equated with high connectivity, this setting incorrectly infers a connection between two highly correlated, yet structurally unconnected nodes. This is another instance of the previously mentioned network effect. When the activity of all nodes is observed, measures like conditional Granger causality should differentiate between direct and indirect coupling, and therefore alleviate the problem. Common input detection requires an underlying network model to estimate the parameters of the input [25], [26], which is at the core of EC estimation methods. Improved techniques, such as partial correlation, have been devised to alleviate this problem, but they also result in symmetric connectivity matrices and tend to give reliable estimates only in sparse networks. A special form of partial correlation was derived for spiking networks and can detect both indirect coupling and also the directionality of connections [27], [28]. In summary, a major shortcoming of FC is the question of how to interpret the observed activity correlations in terms of the effects that caused them independently of structural connectivity (for example, common input).

Answering this question requires model assumptions on both single node dynamics and external inputs to the network, as typically done for EC methods [16]. In other words, the assumed model must be inverted to estimate the connectivity, which is beyond the phenomenological level offered by computing covariances.

Recent work on the nematode *C*.*elegans* has also shown that structural connections alone cannot account for signal propagation and, instead, extrasynaptic mechanisms must be taken into account [29]. Furthermore, neuromodulation is thought to significantly shape the activity of neural circuits, and therefore, SC again might only be the ‘minimal structure’ required for a causal understanding of any neural system [30]. Therefore, a causal understanding of neural network function is not provided solely by computing FC or SC alone. However, when taking a dynamical systems theory perspective, EC or SC in the form of the precise values of the coupling matrix of a given system is the quantity of interest and, if the nodal dynamics and the input to each node are known, the matrix entries are the key to a formal understanding of the system [31].

In general, estimating SC, and EC, in neural systems is hard, both experimentally [32] and theoretically [33]. Recently, an approach called dynamic differential covariance (DDC) has been put forward to infer EC [22]. Methods aiming to infer EC typically make assumptions about the structure of the network or the individual node dynamics before inferring connectivity. DDC assumes that any given time series data is either generated by a linear or a non-linear autonomous dynamical system without time delays and uses this structure to estimate EC, essentially by a series of matrix inversions. The method was shown to reliably estimate the directionality of connections even in the presence of confounder motifs, and to exhibit high noise tolerance, in the sense that individual trajectories could be perturbed by noise without a massive decrease in the accuracy of the algorithm [22]. It also showed good performance when applied to non-stationary data, spiking neural networks and resting-state fMRI data. The validation of the method was carried out in cases when a GT connectivity matrix was known, either by design or by biological knowledge [22].

### 1.3 Estimating effective connectivity without ground truth data

Besides shortcomings of any particular connectivity estimation algorithm, another frequent scenario poses a significant challenge: the GT connectivity of a system might not be known. In the absence of GT connectivity data and without a detailed mechanistic understanding of each node’s internal dynamics, it is an open theoretical challenge to reliably estimate EC structure solely from neural time series recordings. A straightforward first approach is the application of many different estimation algorithms: if after application of these, a common structure, for example a network core [32], becomes apparent, one can be confident that the estimation procedure has reliably estimated at least some elements of the real network structure. Clearly, in the absence of GT connectivity data, merely determining a system’s SC does not suffice to fully comprehend the system dynamics, and hence its EC. However, if the system dynamics can be recreated from an estimated SC, assuming valid node dynamics, it would be a strong indication that the connectivity reflects the underlying physical connections in the system. For most systems, this is a formidable task, which in general does not have a unique solution. Indeed, the task of jointly inferring node dynamics and connectivity structure is in general a hard problem in system identification [34], [35].

Typically, some parametric form of node dynamics (for example, linear dynamics with additive or diffusive coupling) is assumed [17], [36], and the connectivity estimation is carried out in a subsequent step. In neural systems, this problem is often exacerbated by their high dimensionality and non-linearity of the underlying generating dynamical system, resulting in many parameters that can give rise to a particular observed activity data set [37], [38]. In a biological context, scenarios where a GT is lacking are especially relevant when it is hard or even impossible to trace actual physical connections, either because of the lack of transgenic animals or because the network under study is not self-contained so that some of the observed dynamics come from outside the system. In general, this scenario cannot be ruled out in real biological systems where rarely all network nodes can be imaged with high temporal and spatial resolution. Apart from lacking information on GT connectivity, in biological systems, nodal dynamics might also not be known. The reason is that again, the nodal dynamics might not be experimentally accessible in detail. This is especially true for non-mammalian model systems, such as cnidarians, among them jellyfish and Hydra [39], [40], [41], [42]. For example, in the case of jellyfish neurons, it might not be feasible to experimentally determine the electrical properties of spike or transmembrane signal transduction. In this case, one has to resort to either qualitative descriptions of the dynamics or first determine the form of nodal dynamics by specialized estimation methods [43], [44].

Thus, in the absence of GT connectivity data, any single estimated connectivity (EStC) matrix is not really valuable per se, because it cannot be fundamentally validated. This is reminiscent of the scenario where only FC is measured, which does not offer a causal understanding of the system. Hence, if only the number of nodes and their corresponding activities are known, it might not be sufficient to just estimate connectivity. Therefore, in the absence of GT connectivity data, the task of connectivity estimation must be augmented by a regeneration step to obtain knowledge on EC. If the nodal dynamics from this regeneration agrees well with that of the real system, the agreement indicates that our choice of dynamical system and SC faithfully reflects the processes that generated the original time series data. Then, graph theory methods can be used to study the connectivity structure of the biological network of interest [45].

### 1.4 Outline of the paper

In this paper, we are interested in unraveling the network interactions that produce observed node activity time series, which amounts to understanding how the system functions by observing node activity. For a complete understanding, we need to infer the EC of the system. As already explained above, EC can be classically decomposed into SC among the nodes plus a model of the local node dynamics, in the sense that estimating EC always involves model assumptions. Hence, the task can be broken down into two steps: (1) infer the SC and (2) define a node model and use it together with the SC to reconstruct the system dynamics. For (1), the different inference approaches need to be validated against GT connectivity data. For (2), they need to be further validated against the observed node activity patterns. In general, SC is represented by sparse directed weighted graphs [21]. When equipped with a model for the node dynamics, it is possible to simulate the entire underlying system, which opens access to EC. Hence, obtaining EC, on top of SC, will result in a more complete understanding of the neural system under investigation.

Here, we study a method, lagged-cross-correlation (LCC) analysis, to estimate SC in a wide range of systems. Our method does not make assumptions on the network topology or dynamics, and as such is model-free. We first focus on mathematical models of neural networks as exemplified by the Hopf model [46], [47] as a system with known GT connectivity, which features non-linearities and spatiotemporal delays. We show that in sparse networks, LCC-based methods as well as methods combining LCC with the derivative-based method DDC outperform other methods of connectivity estimation, including standard-correlation based and purely derivative-based methods by themselves. We then systematically study the performance of different algorithms in terms of the quality of network reconstruction as the connection density varies. For sparse connectivity, the LCC methods perform best, whereas for denser connectivities they perform on a par with other methods, such as standard correlation-based methods. We show that LCC provides reliable estimates when the number of inputs to a neuron does not exceed 5, and link this to the emergence of one giant connected component. We also comment on the performance of our methods when intrinsic parameters of the Hopf model are changed, revealing both monotonic and non-monotonic dependencies of connectivity estimation performance on these parameters. We then study the performance of LCC and DDC in linear networks without delays, and show that in these networks, in contrast to the results for the Hopf model, DDC outperforms LCC and leads to a near-perfect reconstruction of GT connectivity even in dense and large networks. We provide explanations for these observations. We next ask whether our inferred SC matrices can be used to recreate the observed nodal dynamics of the Hopf model, and hence gain insight into EC, to which the answer is affirmative under strict conditions on the random seed of simulated and recreated data. We compare the quality of the data reconstruction when connectivity matrices estimated with different algorithms are used to recreate the system dynamics. Again, we find that our method outperforms derivative-based methods, because it results in higher trace-to-trace correlations. Using our approach, these correlations were perfect for some of the recreated traces. Finally, we apply our method to biological data from the nematode *C. Elegans*, and show that it performs at least as well as a recently devised computationally more expensive reservoir-based method [48]. We comment on the performance of different methods applied to the dataset.

In addition to providing a systematic validation of different methods in two paradigmatic scenarios, our work constitutes a first important step in tackling the problem of validating connectivity in the absence of GT connectivity data. We propose that combining SC estimation with a recreation simulation can serve as a phenomenological system identification step [49]. Furthermore, this reconstruction step can help to bridge the gap between SC and EC connectivity measures. Indeed, having estimated an SC matrix, it can be used as an input for a forward simulation of a system. If the system dynamics is recreated by this step, then the estimated SC matrix also bears relevance for the EC structure of the system.

## 2 Materials and Methods

### 2.1 Methods for connectivity estimation

#### 2.1.1 Lagged Correlation

Our principal new method, LCC, is based on Fourier transforms. In short, the method estimates the lag at which cross-correlation between two signals is maximized and uses this maximal value of the correlation as an approximation for the strength and directionality of connectivity between two nodes. As such, our method is related to the work by [17], [50], [51], [52], who used lagged covariance or inverse correlation matrices to study the directionality of connections in fMRI experiments and spiking neural networks.

Cross-correlation between two signals *X*(*t*) and *Y*(*t*) at lag τ is defined as a measure of similarity between *X*(*t*) and a time-shifted version of *Y*(*t*), *Y*(*t* + τ). The time shifted version *Y*(*t* + τ) is then used in the integral

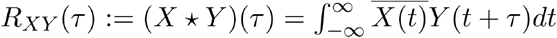

over time to compute cross-correlation between *X* and *Y*. 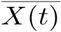denotes complex conjugation of *X*(*t*). This integral calculates how much the signal *X*(*t*) correlates with *Y*(*t* + τ) as a function of the time lag τ.

The integral can be computed directly in the time domain. However, calculating the cross-correlation can be computationally demanding, especially when the signals get longer. Therefore, to efficiently calculate the cross-correlation, the method is using Fast Fourier transformations (FFTs) in multiple steps. The first step is to transform the signals into frequency domain using an FFT:

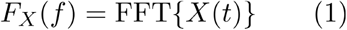

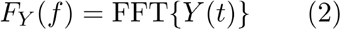

To now compute the cross-spectral density, the method uses the power spectral density by multiplying the Fourier transform of the first time-series with the complex conjugate of the Fourier transform of the second time-series:

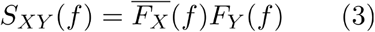

Afterwards, to obtain the cross-correlation in the time domain, the inverse of the Fourier transform is calculated by using the Inverse Fast Fourier Transformation (IFFT)

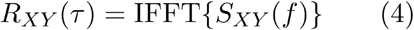

Since an IFFT transforms back from frequency to time domain, the cross-correlation explicitly depends on time. By virtue of the convolution theorem, this inverse is the cross-correlation in the time domain.This calculation outputs a sequence consisting of the cross-correlation of the two time series at different lags. The lag with the highest cross-correlation is determined by finding the index in the sequence at which the correlation is highest, yielding a lag:

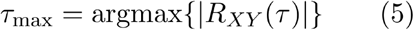

By examining *R*_*XY*(τ)_ and *R*_*YX*(τ)_, the method assesses the directionality of the interaction. If τ_max_ for *R*_*XY*(τ)_ is smaller than for *R*_*YX*(τ)_ it implies a directional influence from *X* to *Y*. Conversely, a smaller τ_max_ for *R*_*YX*(τ)_ implies a directional influence from *X* to *Y*. Equally large values for τ_max_ in both directions with a high correlation suggest a bidirectionality of the influence. Then, the Pearson correlation coefficients are calculated for each neuron pair. With the use of the directionality calculated in the step before, the EStC matrix is filled with the corresponding Pearson correlation coefficients. Alternatively, one could also use lagged covariance, as done by Cecchi et al. (2007), but we found that this does not lead to large differences compared to using standard Pearson correlation. In summary, this method first used the cross-correlation to determine the directionality of connections, and then estimates connectivity with the Pearson correlation coefficient. We remark that the analysis of the ‘best’ lag in covariances has been described in the literature for both fMRI [53], [54], [55], and synthetic data [56].

#### 2.1.2 Lagged Correlation in conjunction with DDC

Another method that was explored was to use the LCC method in conjunction with the DDC algorithm [22]. For this approach, two matrices obtained with the LCC approach and the DDC approach were compared against each other and only values over a certain threshold, typically set at 0.1, that were present in both estimated connectivities were used. The DDC approach was used in its linear form, using a linear estimation non-linearity. At its core, DDC assumes a linear or non-linear dynamical system in the form 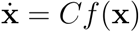, where *f* is either the identity or a non-linear function. This equation is inverted to obtain an estimate for the connectivity matrix *C*. This estimate takes the form of the product of two matrices, namely the covariance between the signals and their derivatives (differential covariance) and the partial covariance matrix (inverse of the covariance matrix). The linear form of the estimator does not need additional assumptions, whereas the non-linear form of the estimator additionally requires the choice of a suitable estimation non-linearity. Under the assumed systems equations, DDC is the least-squares estimator for *C*. For more details, we refer to [22].

#### 2.1.3 Cut methods

For the comparison, the methods described above were also applied to a dataset which was sampled at every 15th time point (i.e., every 15th time point was kept in the time series), resulting in a naive smoothing of the data. This was done to reduce signal-to-noise ratio, which can sometimes be beneficial when working with neuronal data. These methods are marked as “cut” in the comparison matrices.

#### 2.1.4 Thresholded methods

For comparison, some methods were also thresholded after estimation. These are marked as “thresholded” in the comparison matrices. For this, a threshold of 0.1 was chosen, which clamps any value lower than the set threshold to 0. Values above the threshold are not changed. This can be helpful when dealing with sparse connectivity matrices, as these naturally contain many vanishing entries.

#### 2.1.5 Transfer Entropy and Partial Correlation

For the transfer entropy [57] estimated connectivity, the implementation (https://github.com/netsiphd/netrd) from *netrd* [58] was used. This allowed for calculation of the naive transfer entropy (NTE). Partial Correlation was computed using the python package scikit-learn (https://scikit-learn.org/stable/modules/covariance.html). L1 regularization was used for Partial correlation.

### 2.2 Neuron and network models, simulation paradigms

#### 2.2.1 Hopf model

To validate our methods, a Hopf neuron model was implemented using the neurolib library [59] Python package. The Hopf model is a non-linear dynamical system that simulates, through a bifurcation mechanism, the activity of a neuronal network. As such, the Hopf model is not a generic single neuron model, but rather a neural mass model, simulating small patches of neuronal tissue instead of individual neurons [60]. We nevertheless chose this model, because it features complex dynamics and has delays as well as non-linearities, which makes it a good test case for our algorithms. This model is particularly good at capturing the point of transition between stable and oscillatory neuron states, leading to a good representation of the physiological transition of firing states [46]. In this implementation, each neuron’s dynamics are governed by the Stuart-Landau equations, which form the dynamics underlying the Hopf model. Specifically, the two-dimensional dynamics (one variable each for real and imaginary parts of the complex variable whose time evolution is given by the Stuart-Landau equation) of each neuron are described by:

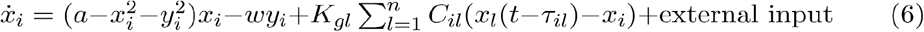

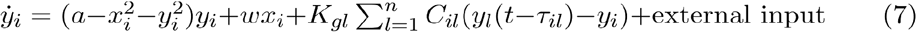

where *x* and *y* represent the real and imaginary parts of the neuron’s state and 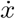 and 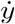 denote a time derivative. The Hopf bifurcation parameter *a* and the natural frequency parameter *w* describe the oscillator. The default values are *a* = 0.25 (non-linear regime [61]) and 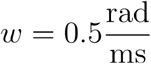, which corresponds to a natural frequency of approximately for each neuron. Thus, the intrinsic oscillation period is approximately 12.5ms. *K*_*gl*_ = 0.6 is the global coupling strength of the model that is multiplied with the connectivity of the current neuron to adjacent neurons from the connectivity matrix *C*. The difference term *x*_*l*_(*t* − τ_*il*_) − *x*_*i*_ (and similarly for *y*) describe the effect of a diffusive coupling scheme. There additionally is a distance-dependent time delay between neurons, defined by a distance matrix *C*_distance_ in the simulator.

This distance matrix is of the same shape as the connectivity matrix and holds values between 0 and 10 at positive values in the connectivity matrix. This in conjunction with a fixed transmission speed yields a time delay in the transmission of the signal.

Concretely, the matrix *C*_distance_ determines transmission delays as follows: whenever C_*ij*_ is 1, the delay between nodes i and j (and also between j and i, if C_*ij*_ is non-zero) is given by

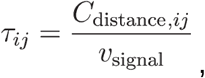

where υ_signal_ is the constant signal transmission speed. We chose 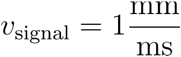, and *C*_distance_ was filled with distances uniformly distributed between 0 and 10, so that the delays τ_*ij*_ where also uniformly distributed from 0 to 10 for our default parameter set. When the delay distribution is changed, we change the distance distribution.

The external inputs are inputs independent of the network dynamics in the form of Ornstein-Uhlenbeck processes (OUPs) *z*, whose stochastic differential equations read 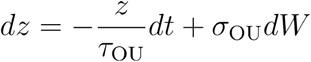, where is a standard Brownian motion (Wiener process), for both *x* and *y*. The default parameters are 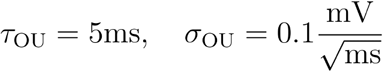. Independent processes were used for each neuron and for each *x*_*i*_ and *y*_*i*_. Each OUP was started from 0. This form of the input is used to represent sensory input or other forms of stimuli that the neurons might experience in a biological context. The system was integrated using the default parameters and settings of *neurolib* [59].

In the context of validating the connectivity estimation method, employing this Hopf model presents a robust approach. First, the dynamics of the model is rich and non-linear. Secondly, by generating a GT before running the model, we can later validate the estimation method by comparing the GT matrix to the estimated matrix. We emphasize that using the Hopf model does not allow for a direct modeling of biophysical processes leading to neuronal firing, for example, via calcium dynamics. Instead, we consider the model to be a good candidate for dynamics with spatiotemporal delays and noise, which strikes a balance between computational complexity and dynamical expressivity.

#### 2.2.2 Linear model

In order to test our algorithms on a standard neuron model without non-linearities or delays (both of which are present in the Hopf model), we resort to the model used by [22], which is given by a single linear equation per node:

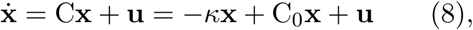

where **x** =[*x*_1,_ *x*_2_,…, *x*_*N*_]^⊤^ is the vector of the state variables *x*_i_, **u** ~ 𝒩 (0, σ^2^) is Gaussian white noise with standard deviation σ and mean 0. Thus, Eq. 8 is a system of coupled Ornstein-Uhlenbeck processes. C is the GT connectivity matrix, in analogy to Eqs. 6 and 7. After the second equal sign, we single out the value κ of the diagonal entries of C together with a matrix C_0_ that has zeros on the diagonal.

Thus, C = − κ𝕀 + *C*_0_, where 𝕀 is the identity matrix. For numerical stability, κ was chosen such that 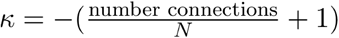. The entries of C_0_ were set to 1, and their sum hence equals the number of connections in the network. Like in Chen et al., all nodes had decaying dynamics, but they were linked by excitatory, instead of inhibitory connections. In contrast to Chen et al., we do not consider observational noise acting on **x**. The system given by Eq. 8 is integrated using an Euler-Maruyama scheme with a timestep of 0.01 s and the system is simulated for 1000 s [22], [59], which results in a simulation length of 100000 time steps.

#### 2.2.3 Network models and data generation

The neural network model was structured as a network consisting of *N* =(10,100,250) neurons using the neuronal models described in Sec. 2.2. The connections between these neurons were chosen based on a pairwise probability p. For method validation, this parameter was initially set to 0.1 and then increased to 1.0 for *N* =10. For *N* =100, was varied from 0.01 to 0.1, and for *N* =250, it varied from 0.004 to 0.04. This entails that the product *pN* is constant for different *N*, which makes the networks more comparable. Our application of this method resulted in a directed, non-symmetric graph (directed random graph): if there is a connection from node i to j, it does not mean that there also is a connection from j to i. *p* is the probability that two given neurons in the network will be connected, but only in one direction, so that the probability of bidirectional connections in our adjacency matrix is low, but increases with increasing p. The total number of connections is given by *pN*(*N* − 1) where *N* is the number of neurons. In the Hopf model, the diagonal is also filled with zeros, such that there exists no connection from a neuron to itself. Such sparse connectivity was an explicit choice, aimed to emulate sparse connectivity patterns observed in biological neural networks. For the Hopf model, the generated datasets consisted of a time series of 200000 time steps. All statistics were computed on the entire length of these time series, which are assumed to be stationary. If not mentioned otherwise, *M* =100 realizations with independent random sees were performed per parameter value.

#### 2.2.4 Forward simulation

To validate the accuracy of the estimated connectivities, we used the same method s as described in Sec. 2.1. First, we estimated the connectivity matrix as described in Sec. 2.1. Then, the Hopf model (Eqs. 6 and 7) was simulated forward in time with the EStC matrix. Applied with the same initial parameters as the original data, this approach was used to recreate the time series data. Furthemore, the seed for the Ornstein-Uhlenbeck processes in Eqs. 6 and 7 had to be set exactly the same, as otherwise the background noise and corresponding oscillation would not match up.

## 3 Results

### 3.1 Comparison of methods for connectivity estimation in the Hopf model

We started to use the LCC method (Sec. 2.1.1) on small networks (*N* = 10) of the Hopf model (Sec. 2.2) to estimate a known GT connectivity. The GT connectivity was chosen to be a sparse matrix with p=10% connectivity. The method demonstrated a remarkably high accuracy in estimating the underlying connectivity of the Hopf neuron model using the proposed LCC approach (Fig. 1). The Pearson correlation coefficient between the EStC and GT connectivities was calculated to be 0.97 (Fig. 1B), which indicates a near-perfect agreement of GT and EStC. For the same data, the DDC algorithm showed a corresponding correlation coefficient of only 0.52 (Fig. 1C).

**Figure 1:**
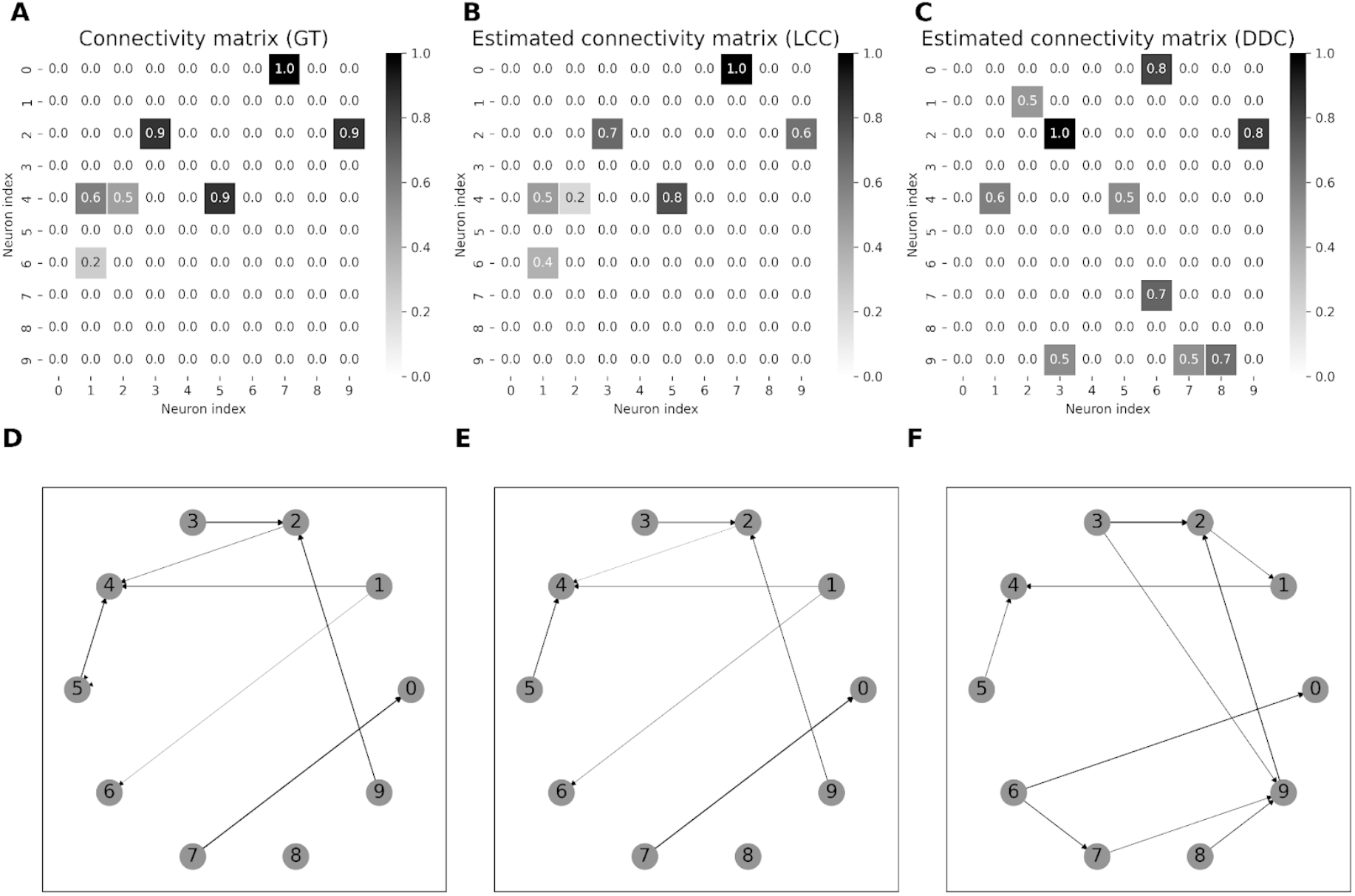
Comparison of GT and EStC connectivity matrices in a Hopf Model using the thresholded LCC method, connection probability p = 0.1. Panel A shows the original GT connectivity matrix used to generate time-series data with a Hopf model. Panel B shows the EStC matrix using the LCC method (Sec. 2.1.1). Nearly perfect agreement with the GT matrix is observed. The Pearson correlation coefficient is 0.97 between the matrices. Panel C shows the EStC matrix using the DDC method. This method finds some of the right connections, but the connection strength is off-target, and there are many false positives. This leads to a lower correlation (0.52) between the EStC matrix computed with DDC and the GT connectivity matrix (panel A). Due to rounding to one decimal place, 2 values are not shown in the GT connectivity matrix, this leads to 7 (instead of 9) visible connections in the GT connectivity matrix. Panels D-F show the corresponding network graphs for the GT connectivity (Panel D) and the two estimated graphs for LCC (Panel E) and DDC (Panel F). Two connections in Panel D are hardly visible because of their small numerical value.

Using the LCC method, we could sometimes even estimate connectivities that near-perfectly recovered the GT connectivity (meaning that both the presence or absence and in case of a connection also its strength was correctly estimated), as shown in Fig. 1 A and B.

Which method performs best for this synthetic Hopf model data set? To answer this question, we systematically studied the performance of different algorithms in small networks of fixed size and increasing connectivity (Fig. 2). Performance here was quantified with the Pearson correlation coefficient between GT connectivity matrices and their reconstructed EStC counterparts. To summarize, the best method for sparse networks is LCC in conjunction with DDC, but LCC on its own also performs well.

**Figure 2:**
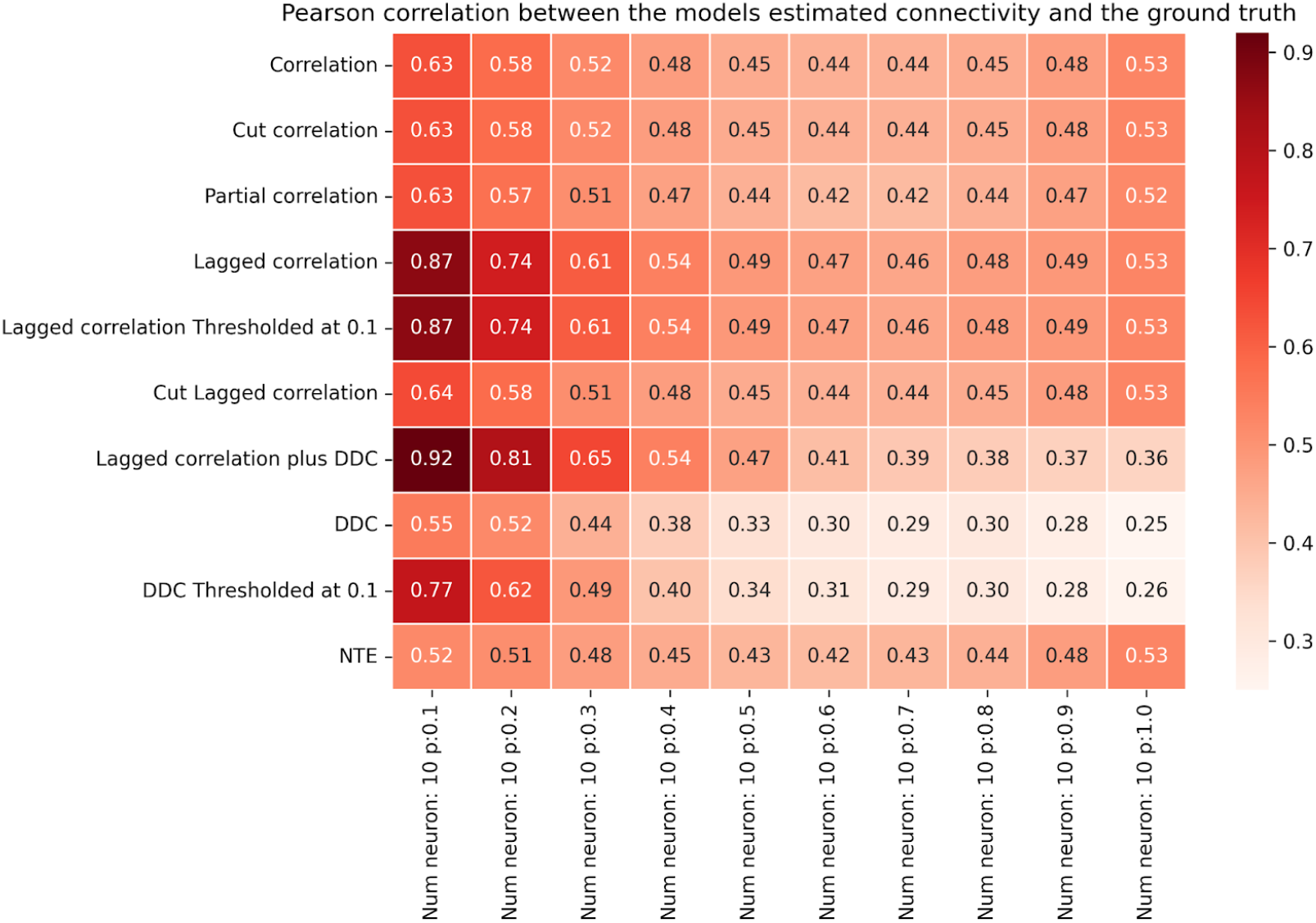
Performance of different methods for for *N* = 10 the Hopf model. Pearson correlation coefficients between EStC matrices estimated using different methods and the GT matrix of the given model parameters for a Hopf model. The number of neurons is set to *N* = 10, while the connection probability *p* increases from 0.1 to 1.0. For small *p* (*p* ≤ 0.5), LCC in conjunction with the DDC algorithm yielded the highest correlation out of all the methods used (row 7). At higher values of *p* correlation-based methods (rows 1 and 2) and cut LCC (row 6) perform best. LCC and thresholded LCC (rows 4 and 5) perform equally well. All LCC and correlation-based methods perform better than DDC-based methods (rows 8 and 9). NTE and Partial Correlation (rows 10 and 3) show fewer changes in performance as *p* is changed, and perform worse than LCC-based methods at low *p* and equally well at higher *p*, where they also perform better than DDC-based methods. The value per cell is calculated by taking the mean of the correlation in 100 simulations rounded to 2 decimal places.

DDC-based methods show lower performance than LCC-based ones, especially at higher connection densities. Standard methods, including Partial Correlation and NTE, perform better than DDC, but worse than LCC. The performance of all methods initially decreases sharply with connection density. LCC-based methods show high performance until the average number of neighbors reaches 4-5.

In more detail, our first result is that the newly proposed LCC method in conjunction with the DDC algorithm (Sec. 2.1.2) showed the best performance for small values of *p* (Fig. 2, row 7). By just using the LCC method (Fig. 2, row 4), the correlation coefficient was not decreasing as rapidly, compared to DDC, with increasing *p*. For all values of *p*, DDC alone and also its thresholded variant (Fig. 2, row 8 and 9) did not perform well compared to LCC-based methods. Moreover, the LCC method performed nearly equally as well on its own (Fig. 2, row 4) as in conjunction with DDC (Fig. 2, row 7), while cut LCC (Fig. 2, row 6) and methods using standard correlation (Fig. 2, rows 1 and 2) showed comparable (lower) performance than LCC-based methods for higher (lower) values of *p*. The agreement with the GT connectivity, however, was much lower than for small values of *p* for all methods. The thresholded DDC method (Fig. 2, row 9) generally yielded a higher correlation for larger values of *p* compared to DDC alone (Fig. 2, row 8), as this method produces true zeros in the EStC matrix leading to a higher correlation especially in the simulations with sparse connectivity.

LCC thresholded (Fig. 2, row 5) performed as well as LCC (Fig. 2, row 4) for all values of *p*. Again, the methods that did not use DDC worked better than those using DDC for larger values of *p*. As a comparison with more traditional methods for estimating directed connectivity, we also computed NTE (Fig. 2, row 10) and Partial Correlation (Fig. 2, row 3). We found that NTE performs worse than LCC for small *p*, but increasingly better at large *p*, where it performs on par with LCC. However, NTE still performs slightly worse than standard correlation-based measures (Fig. 2, rows 1 and 2) and also than cut LCC (Fig. 2, row 6). For nearly all values of *p*, NTE performs better than DDC (Fig. 2, row 8). Thresholded DDC (Fig. 2, row 9) initially performs better than NTE, but then its performance drops below that of NTE. The results for Partial Correlation are similar: initially, the performance is lower than for LCC and comparable to standard correlation-based measures, but then the performance catches up and all three method classes (LCC, correlation, including Partial Correlation, and NTE) are comparable for larger *p*. All three method classes perform better than DDC, especially at higher values of *p*.

How do these results depend on the parameter values of the Hopf model, in particular, on the parameter *a* controlling the dynamical regime, and on the delay distributions (see Sec. 2.2.1 for a definition of the parameters)? For the standard set of parameters, we chose uniformly distributed delay between 0 and 10 ms. We computed receiver-operating characteristics (ROCs) in the SI, Figs. S1 and S2 for fixed delays of 100 ms (Fig. S1) and 10 ms (Fig. S2). The parameter changing in these figures is the threshold for binarization, so that only the binary connectivity (irrespective of the weights) is considered. We find that for small delays, the thresholded LCC method performs best (cf. Fig. 1), whereas for large delays, thresholded DDC performs best. The best threshold value is close to 0.1 for all methods.

Next, we determined how the Hopf bifurcation parameter *a* influences the results. For distributed delays as in Fig. 2, results are shown in Fig. S3. The performance of all methods drops as *a* is increased from initially negative values beyond the bifurcation point (*a* = 0) to positive values, which corresponds to a transition from a linear to a non-linear regime [61]. Increasing the delay in Fig. S4 does not influence the trend with respect to *a*. For large positive *a*, the thresholded DDC method performs best (Fig. S4). We observe that LCC and LCC plus DDC perform better than for larger delays, and on par with thresholded DDC and LCC.

The influence of the noise parameters σ_OU_ and τ_OU_ is studied in Figs. S5 and S6, respectively. We observe a non-monotonic dependence on the estimation performance: the Pearson correlation between GT and EStC matrices peaks at an intermediate value for the noise parameters. An intuitive explanation for this is that the dynamics for small noise parameters is not rich enough, whereas for larger values of the noise parameters, it shows too much randomness. This interesting observation will be studied in more detail elsewhere.

A systematic parameter sweep (SI, Figs. S7-S12) concludes our study of the parameter dependence of the estimation performance. In these figures, the Hopf bifurcation parameter *a* is fixed in any one figure, whereas the fixed delay changes. In the non-linear regime (Fig. S7), LCC plus DDC performs best, followed by LCC thresholded and LCC and DDC thresholded. For larger delays, DDC thresholded performs best, followed by standard DDC and LCC thresholded. This trend is similar for more negative values of *a* (Fig. S11, note again that estimation performance decreases with increasing *a*), with DDC thresholded being the best method for larger delays. LCC and its thresholded or cut variants, however, do not perform much worse and are still in the top three methods for all values of *a* and larger delays. To summarize, LCC plus DDC is the preferred method for small delays, whereas DDC thresholded is the best method for larger delays, across all values of *a* (Fig. S12). LCC and LCC thresholded are not much worse than LCC plus DDC for small delays. Standard DDC does not perform on par with these methods for any delay.

Summarizing, we have shown that our comparatively simple method and its variants can successfully reconstruct model connectivity from dynamics in a non-linear model with spatiotemporal delays. In general, all methods perform worse when *p* is increased. We find that *pN* ≤ 5 is the approximate threshold where the performance of our LCC algorithm drops rapidly (Fig. 2, column 5 vs. column 6). This also holds true for larger networks (Figs. 3-4, column 5 vs. column 6).The threshold value for *pN*, after which performance drops below 0.5, is slightly higher for *N* = 250 than for *N* = 100 and *N* = 10, with a value of *pN* ≤ 6. A connected component analysis (SI, Figs.S13 and S14) revealed that, for larger networks the singular connected component occurs at a higher *pN*, and hence, we expect that performance in these networks does not decay as early as in smaller networks as the connectivity is increased. In summary, our methods work best for sparse networks, presumably such as those found in the cnidarian *Hydra* [41], the nematode *C. Elegans* or jellyfish [42]. In evolutionary higher animals like zebrafish, drosophila, mice and rats, it can well be the case that networks at the cellular level exhibit less sparse connectivity [8], [62].

**Figure 3:**
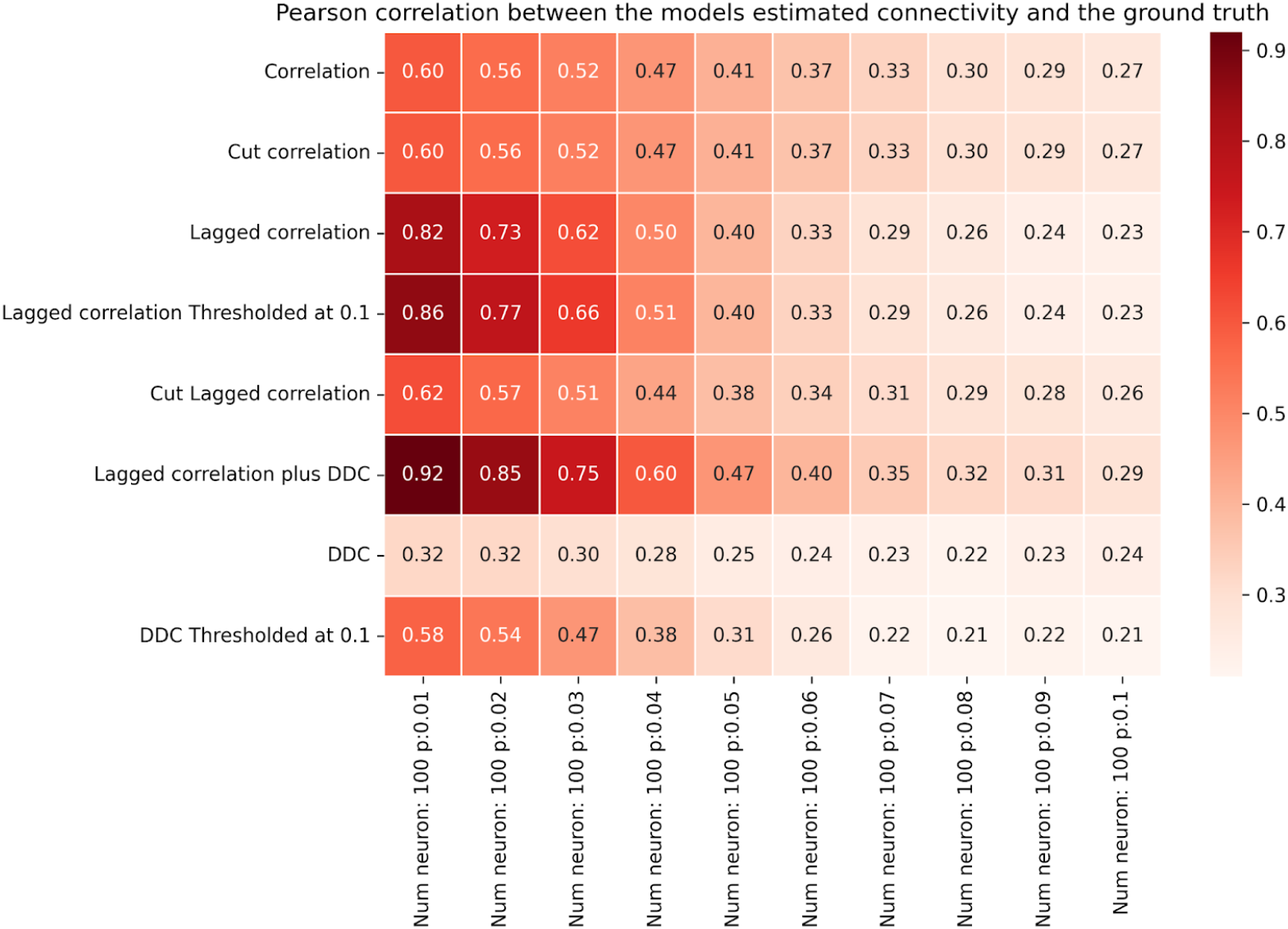
Performance of different methods for *N =*100 for the Hopf model. Pearson correlation coefficients between EStC matrices estimated using different methods and the GT matrix of the given model parameters for a Hopf model. The number of neurons is set to *N =*100, while the connection probability *p* increases from 0.01 to 0.1.

How large can the networks be for LCC to show successful performance? In light of the results shown in Fig. 2, we expect that for larger networks, the performance will drop when *p* is fixed. To facilitate comparison, we keep *pN* constant and show results for larger networks in Figs. 3 and 4. Indeed, as in Fig. 2, the performance of our best algorithm (LCC in conjunction with DDC) drops when *pN* > 5. This happens when moving from column 5 to 6 in Figs. 2-4. Hence, the performance of our algorithms is approximately independent of network size (with a suitable adjustment of p, so that *pN* is constant), and only the average number of inputs to a neuron, given by *pN*, determines performance of the algorithms. This is comparable, both qualitatively and quantitatively, to other studies [51], [63], [64], [65], [66], [67], [68]. It is also theoretically expected [63]. In SI, Fig. S13 and Fig. S14, we confirm that as *pN* is increased, the number of both weakly (Fig. S13) and strongly (Fig. S14) connected components of the GT connectivity matrix decreases. As *pN* approaches 5, all networks tend to have only one weakly connected component, whereas they still tend to have more than one strongly connected component at this value of *pN* for all *N*. The decrease in the number of connected components is stronger for smaller networks and reaches 1 more quickly for *N* = 10 than for *N* = 100 and *N* = 250, as expected. As expected, the onset of decay performance is not as early for *N* = 250 (Fig. 4) as it is for *N* = 10 and *N* = 100 (Figs. 2 and 3).

**Figure 4:**
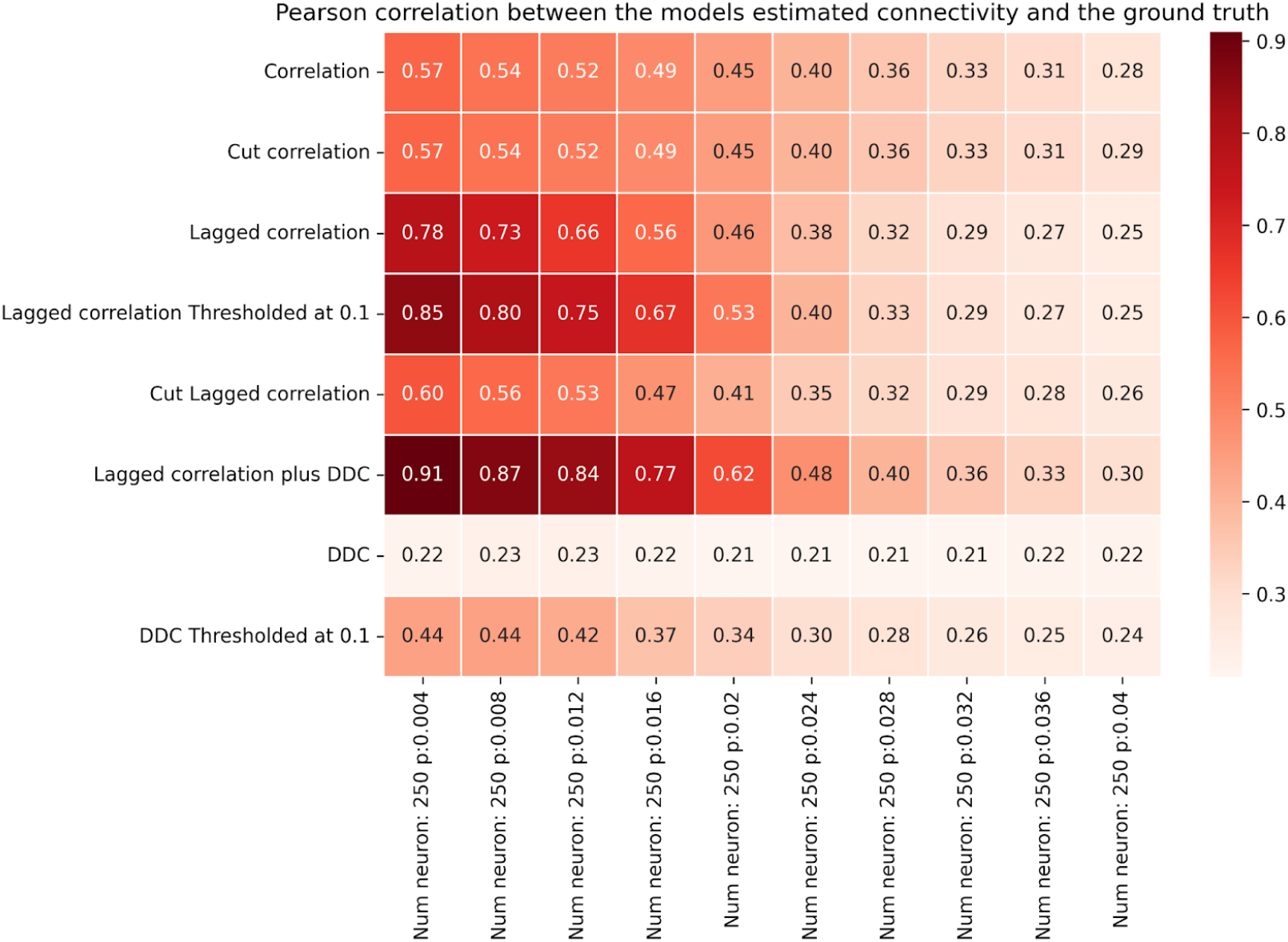
Performance of different methods for *N =*250 for the Hopf model. Pearson correlation coefficients between EStC matrices estimated using different methods and the GT matrix of the given model parameters for a Hopf model. The number of neurons is set to *N =*250, while the connection probability *p* increases from 0.004 to 0.04.

### 3.2 Comparison of methods for connectivity estimation in the linear model

We next asked how our methods compare in networks where the node dynamics are given by linear stochastic differential equations (Eq. 8). With the assumption of linear dynamics, we expect that DDC will result in a near-perfect reconstruction of the GT connectivity when sufficiently long trajectories are considered. The reason is that DDC is the least-squares error estimator for the linear system given by Eq. 8 [22]. Hence, for infinitely long trajectories, we expect a perfect reconstruction of the GT connectivity. For one specific case, we indeed confirmed that the performance of LCC increases as we consider longer trajectories (SI, Fig. S15), which is also expected theoretically [51], [63], [64].

For small networks, the high performance of DDC is confirmed in Fig. 5: DDC (Fig. 5, row 8) and especially its thresholded variant (Fig. 5, row 9) show high performance nearly independently of *p*. Methods involving LCC (Fig. 5, rows 4-7) show lower performance than DDC for all values of *p*, but especially for larger values of *p*. Given that for *p* = 1.0, the networks are fully connected, this is due to false negatives:

**Figure 5:**
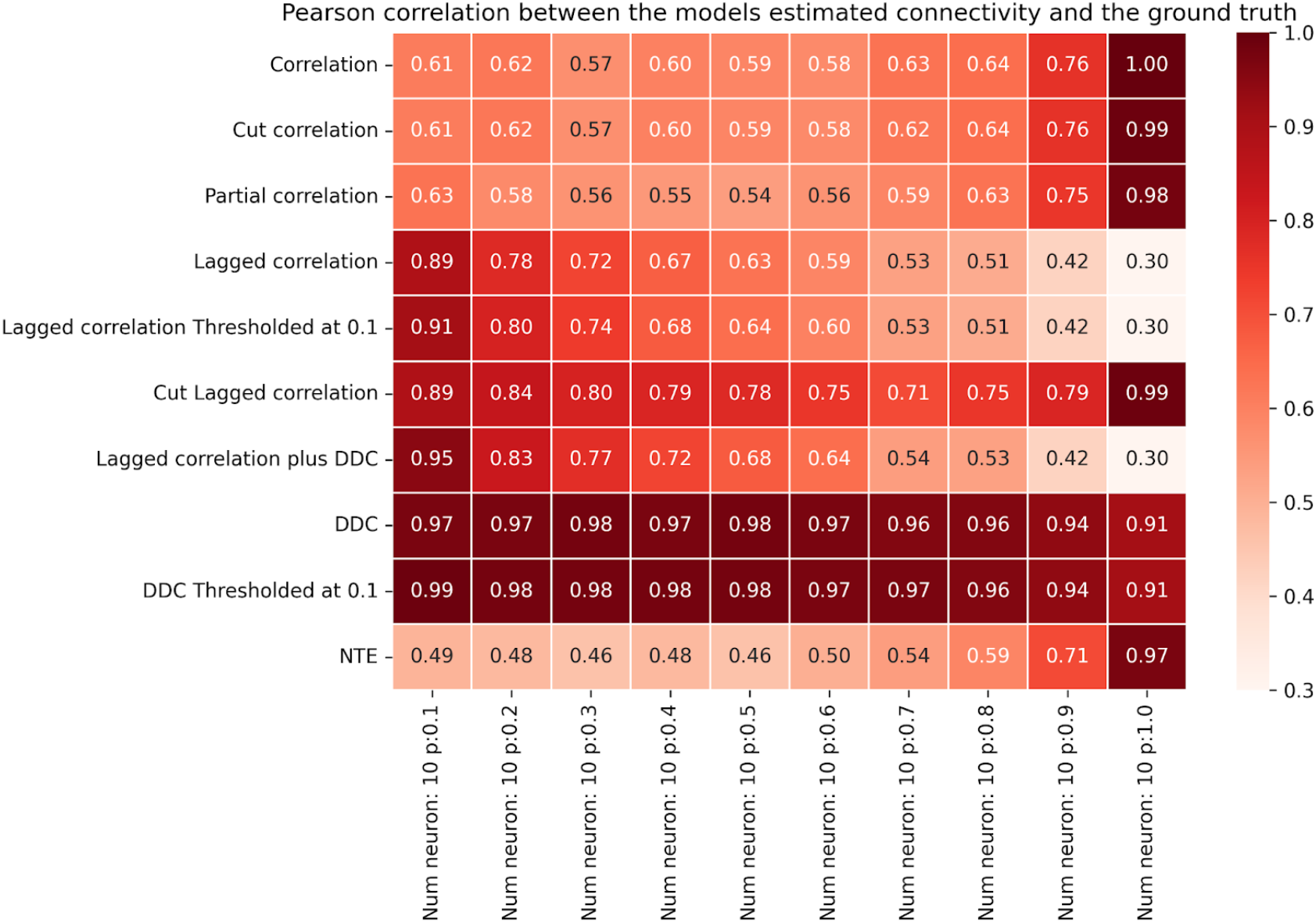
Performance of different models for *N* = 10 for the linear model. Figure layout similar to Fig. 2, but for the linear neuron model (Eq.8).

LCC-based methods miss many connections that are however present in the system. Correlation-based methods (Fig. 5, row 1, 2 and 3) show first a slight decrease, then an increase of performance as *p* is increased. This is because for large *p*, in addition to being less sparse, the matrices are also increasingly symmetric. Cut LCC (Fig. 5, row 6) surprisingly shows a similar behavior to correlation-based measures for larger *p*, and outperforms them for smaller *p*, which shows that in the absence of delays, naively smoothing the time series might be advantageous for LCC. The performance of NTE (Fig. 5, row 10) initially does not increase with connection probability and initially shows lower performance than DDC-, LCC- and correlation-based methods. With an increase of *p* past *p* = 0.5, NTE performs on par with DDC-based and correlation-based methods, likely due to more symmetric connections in the network.

These general trends continue in larger networks (Figs. 6 for *N* = 100 and 7 for *N* = 250), where again, DDC-based methods outperform all other methods and show high performance that is nearly independent of *p*. Especially for *N* = 250, the thresholded DDC variant shows a higher performance than DDC without thresholding (Fig. 7, rows 7 and 8). In this case, DDC also slightly increases its performance as *p* is increased, which means that the method can give rise to false positives at small values of *p*. Cut LCC (Figs. 6 and 7, row 5) initially shows better performance than correlation-based methods. For larger *p*, both methods show similar performance. Overall, however, the performance of LCC-based methods is lower than that of DDC-based methods.

**Figure 6:**
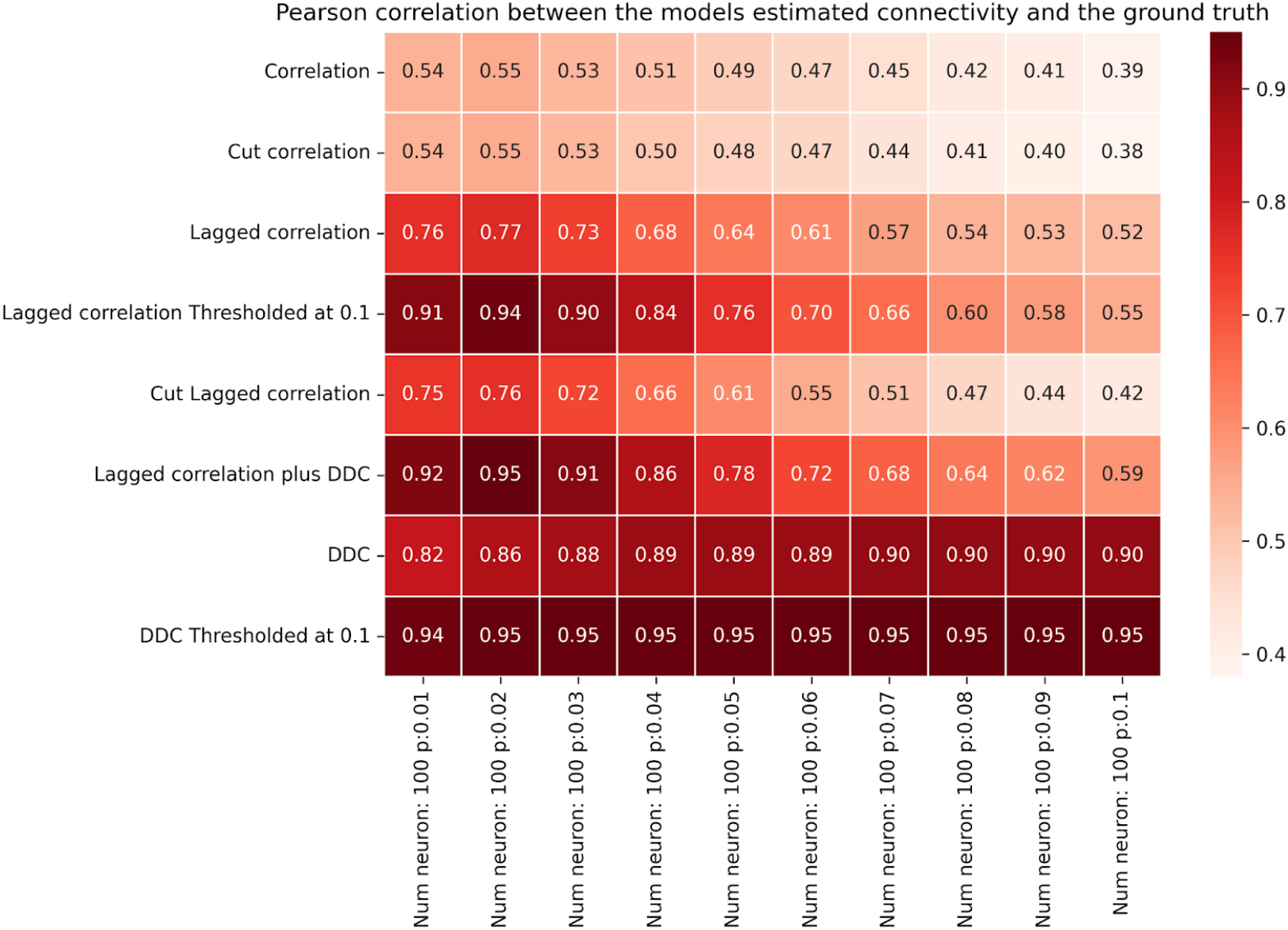
Performance of different models for *N* = 100 for the linear model. Figure layout similar to Fig.2, but for the linear neuron model (Eq. 8).

**Figure 7:**
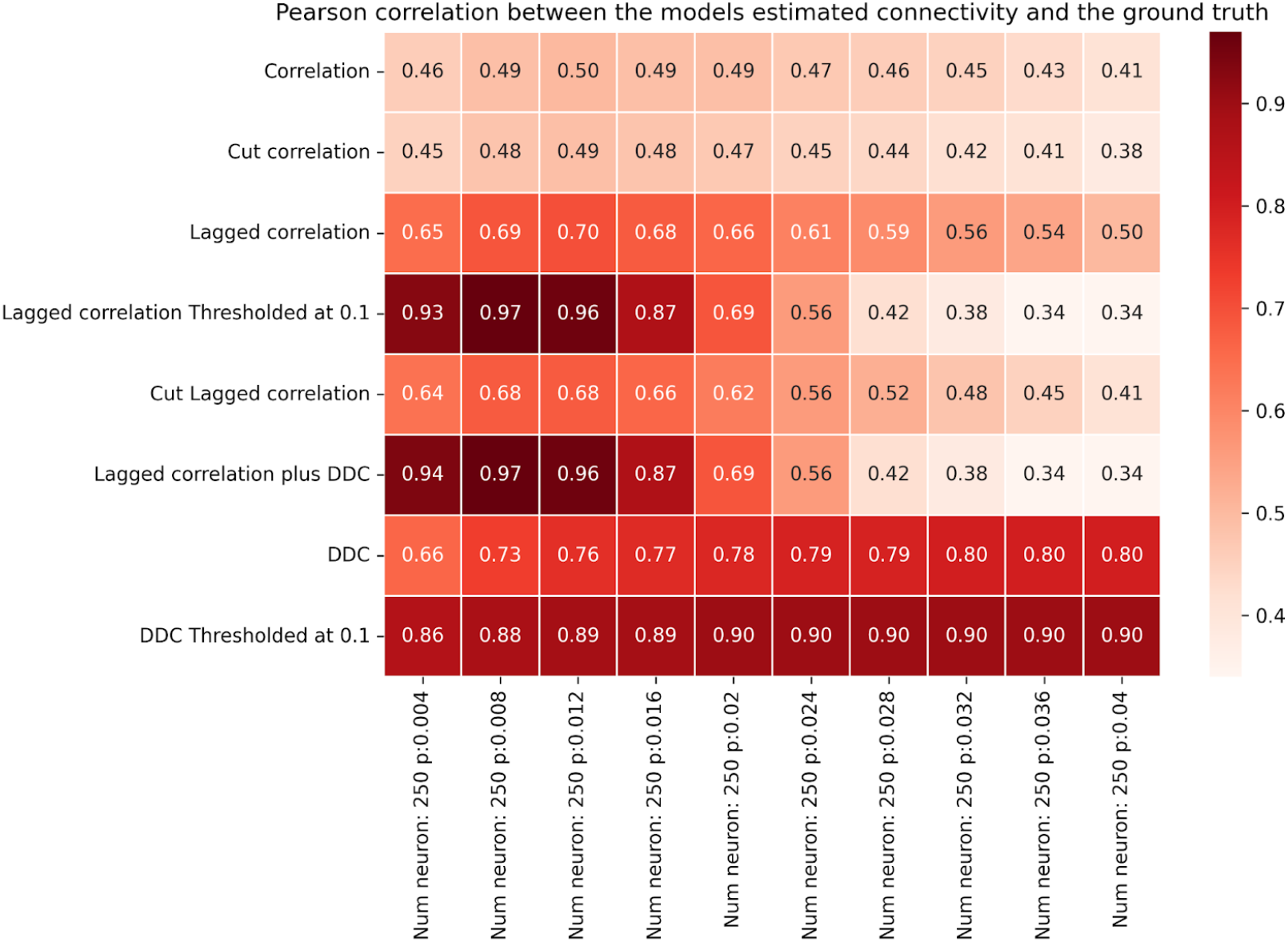
Performance of different models for *N* = 250 for the linear model. Figure layout similar to Fig. 2, but for the linear neuron model (Eq.8).

In summary, we find that, as expected, DDC is a near-perfect estimator for linear systems under the influence of noise. Our LCC method is inferior to DDC in this case.

In Fig. S16, we verify that the near-perfect estimation performance of DDC for linear dynamics deteriorates as fewer time points are used for the estimation (‘cut methods’). This is especially true for cut LCC plus DDC. The performance of cut LCC thresholded degrades most gracefully, followed by cut LCC and cut correlation. Cut DDC shows lower performance than LCC-based methods. Thus, the results presented in Figs. 5-7 are an ideal case, in which the uncut and thus non-subsampled signal is used for estimation. This is in line with theoretical results for estimating connectivity in linear OUP-like systems [17]. For the Hopf model, we observe similar results (Figs. S17-S18).

Due to the non-delayed dynamics of the linear model, the LCC algorithm struggles at increasing *p* to correctly identify the directionality of the data, which is present in the GT coupling matrix. This leads to an increase in false negative rates as the wrong directionality might be inferred. Mechanistically, we find that LCC misses bidirectional connections that are present in particular for larger connection probabilities. It can also wrongly infer the directionality for small connection probabilities as no delays reflecting directed connectivity are present in the data. Especially for larger connection probabilities, the correlation between all nodes rises without the use of a delay. This further decreases the ability of LCC to correctly determine the directionality or even the mere presence of a connection. This rapid decrease happens especially when only one connected component is left, meaning each neuron *a* has a path to neuron *b* and vice versa. Thus, surprisingly, the presence of delays, often assumed to be a complicated dynamical feature, is so-to-speak beneficial for the performance of our LCC method, and detrimental for DDC-based methods.

We finally wondered whether NTE could outperform LCC at a comparable computational cost. To quantify this, we performed a simulation of the linear model (Eq. 8) for increasing *N* and estimated connectivity using either LCC or NTE. The results are shown in the SI, Fig. S19. Whereas for LCC, computational cost as measured by execution time merely increases from 0.4s for *N* = 10 neurons to approx. 60s for *N* = 10 neurons, it increases from approx. 34s to nearly 8000s for NTE. For each individual N, execution time for NTE is two orders of magnitude larger than for LCC, while the accuracy for LCC is higher than for NTE, albeit only by a small margin (inset in Fig. S19, compare also to Fig. 5 for *p* = 0.1). We conclude that for small sparse networks, LCC is the preferred method as it yields higher accuracy at a much lower computational cost. For computational tractability, we did not include computations for NTE and Partial Correlations for the larger networks shown in Figs. 3,4,6 and 7.

### 3.3 Regeneration of node dynamics with different EStC matrices

We next asked the question whether connectivity matrices estimated with our methods can be used to recreate the time series of each node (see Sec. 2.2.1 and 2.2.4 for details). Fig. 8 presents a comprehensive comparison between two distinct methods for estimating neuronal connectivity: LCC (traces labeled ‘simulated’) and DDC (traces labeled ‘dcov simulation)’. The analysis is conducted on simulated data from two neurons. Each plot illustrates three trajectories: the original data (‘original’, GT simulation), the simulated data reconstructed using the LCC method (‘simulated’), and the simulated data reconstructed via the DDC approach (‘dcov simulation’). The GT matrix is shown in Fig. 1A. Fig. 8A shows neuron activity traces where the LCC method correctly identified all the connections as 0, therefore yielding almost perfectly similar traces. This is a special case for our sparse connectivities, as some neurons are not connected to any others (cf. row for neuron 1 in Fig. 1A) and if this is estimated successfully, the correlation is 1 as all the activity is derived from the underlying oscillation (Eq. 6 and 7). In other words, the coupling matrix does not influence the dynamics of this particular neuron 1. The regenerated traces using DDC results in a slightly lower value for the correlation (0.94). Panel B shows neuron activity traces for a neuron in the model where the LCC method estimated a very similar connectivity compared to the GT connectivity This results in a trace-trace correlation equal to 0.97. Using DDC to determine the EStC, the trace-trace correlation drops to 0.79. Our regeneration approach can be considered successful whenever the trace-trace correlations averaged over all traces exceed a certain threshold, for example, 0.8. Concretely, the correlation of the simulated data using LCC with the original GT data is 0.91 over all the 10 neurons, while the correlation using DDC only reaches 0.8 (SI, Fig. S20). In Fig. S20, we show all ten traces of the simulation shown in Fig. 8 (where original and regeneration simulation have the same seed), showing that the correlation between original and LCC-regeneration simulation are higher than those between original and ‘dcov’-regeneration for all ten neurons.

**Figure 8:**
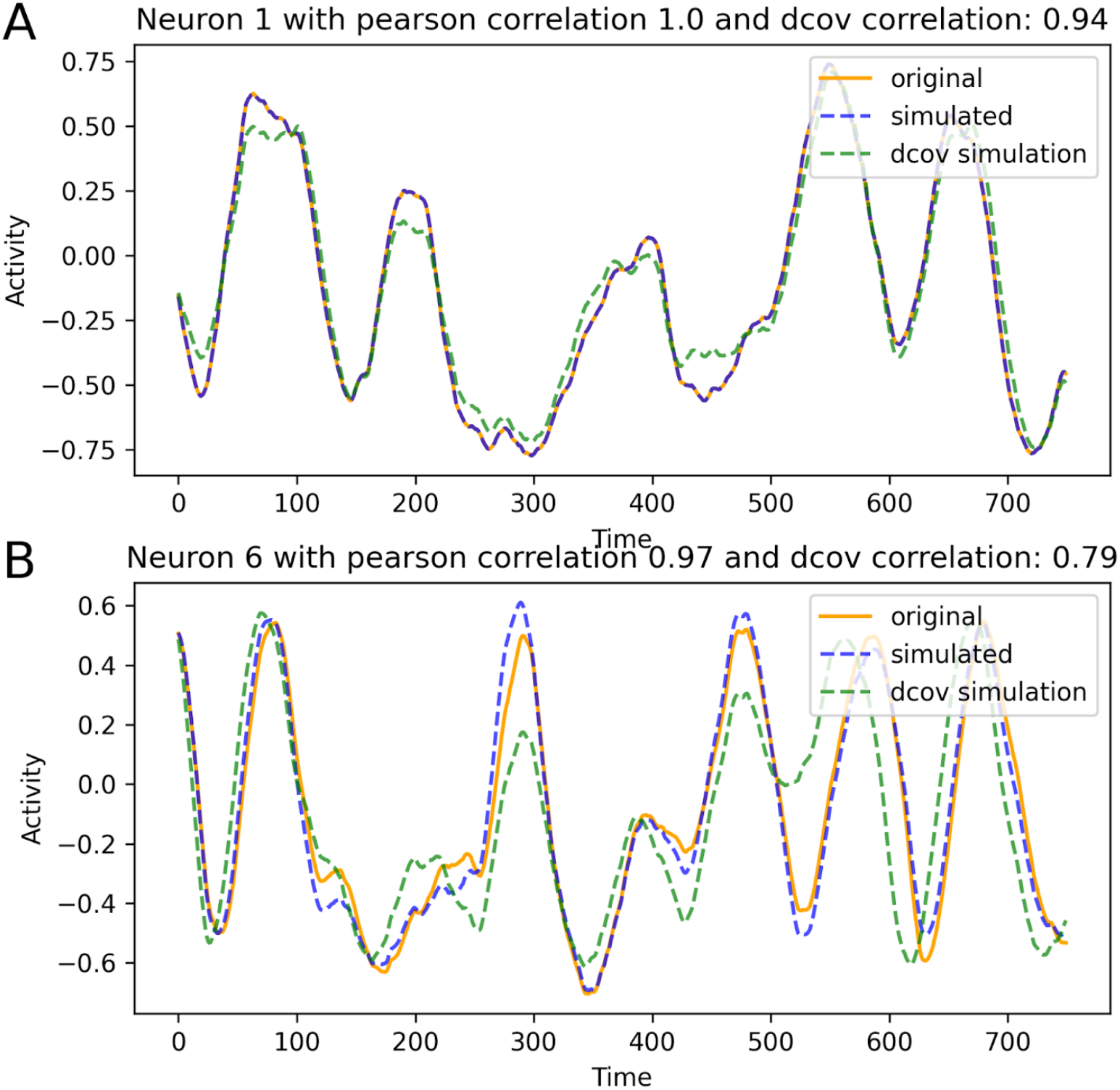
Comparison of the simulated data using the GT, LCC-estimated connectivity and DDC-estimated connectivity for a network of 10 neurons with p=0.1. Orange: Original simulation. Blue: Regeneration of the dynamics using LCC to obtain EC. Green: Regeneration of the dynamics using DDC to obtain EC. Panel A: Results for neuron 1. Panel B: Results for neuron 6. The real part of the corresponding Stuart-Landau equation (Eqs. 6 and 7) is here plotted as activity. The same random seed for the stochastic input to each node’s internal dynamics was used in original and both recreated trajectories.

The main reason for the lower correlation when using DDC is that the EStC matrices are less sparse: incorrectly identified connections of a neuron can affect other neurons indirectly connected with it. A typical scenario is that of secondary connected neurons. If the EStC matrix wrongly results in a connection between neuron 1 and neuron 2, and neuron 2 is correctly connected to neuron 3, then neuron 1 wrongly influences the activity of neuron 3. This lowers the correlation of the reconstructed activity of neuron 3 with the GT activity. Thus, false positives in the connectivity matrix result in low correlations between reconstructed and GT dynamics. In summary, we have shown that if the EStC is close to the GT matrix, then the node dynamics can be accurately recreated in a forward simulation of the system. For good quantitative agreement of the regenerated traces with the original traces, evidenced by high trace-trace correlations, the EStC matrix must nearly perfectly resemble the GT matrix. Small differences in the two matrices result in larger differences between the traces. In the SI, Fig. S21, we show results similar to Fig. 8, but when the seed used for original and regeneration simulation differ. This results in lower correlations between original and the LCC regeneration simulation, which are now comparable to the ‘dcov’ simulation. Thus, in this case, not using the same seed has a similar effect as using a less accurately estimated EStC matrix.

The noise parameters σ_OU_ and τ_OU_ also influence the regeneration performance (SI, Figs. S22 and S23). Regeneration performance, as measured by the average correlation between original and regenerated traces, peaks at intermediate values for the noise parameters. The reason is similar to the non-monotonicity observed for estimation performance (Sec. 3.1 above). For small values for the noise parameters, the dynamics is not rich enough, whereas for large values, the noise is too strong, leading to amplification of small errors in the EStC.

### 3.4 Application to *C*.*elegans* data

Finally, to test our methods using empirical biological data, we applied the LCC method to a dataset of the 8 most active motor neurons of the nematode *C. Elegans* [69]. This is the same data as used by the recent study [48], in which a reservoir computing approach was used to infer connectivity. Applying our LCC method to the same data results in a higher count of true positive values (21 compared to 14) for the same amount of false positives (4 FP), as shown in Fig. 9. Note that the GT data in this case is binary, so a suitable binarization threshold is needed after application of the LCC method.

**Figure 9:**
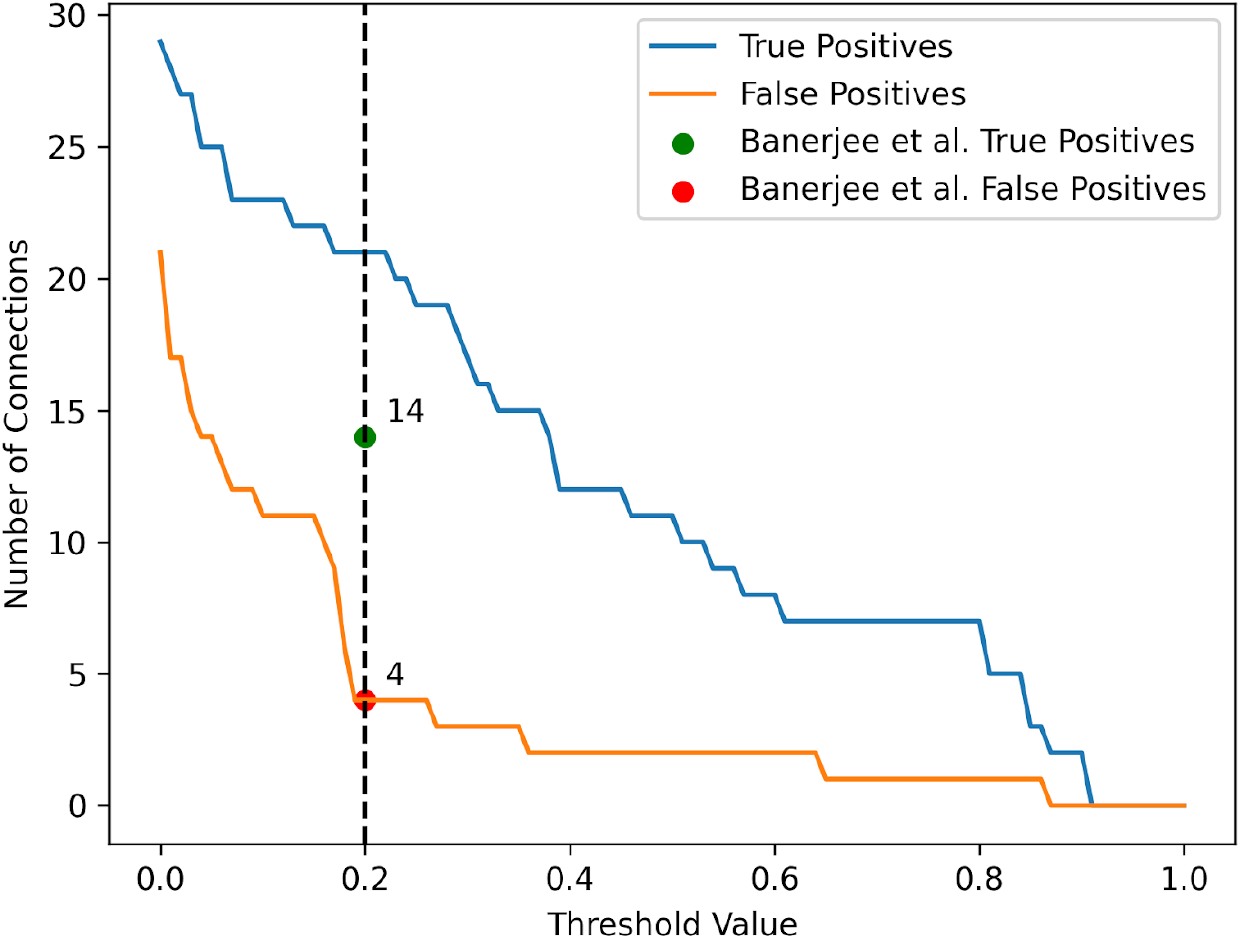
Performance analysis of LCC method binarization thresholds on true and false positive connections in C. Elegans time series data. The figure depicts the relationship between the cut threshold, seen on the x-axis and the corresponding counts of TPs and FPs. A threshold value of 0.2 (black dashed line) was chosen to have comparability against the methods proposed by Banerjee et al. [48].

The number of true positives (TPs) and false positives (FPs) as a function of the binarization threshold value is shown in Fig. 9. The threshold clamps all the values above it to 1 and everything below to 0 to obtain a binary connectivity matrix. Considering the slope of the curve for FPs in Fig. 9, another way to choose the threshold value is given by placing it just before the number of FPs increases sharply, which is just below 0.2. Concretely, in decreasing the threshold from 0.2 to 0, FP increases from 4 to 21, whereas TP increases much less steeply from 21 to 29. The green and red dots in the plot show the TP (14) and FP (4) values from the reservoir computing shown in the recent study [48].

This shows that the suggested method using LCC does not only perform well when applied to simulated Hopf neuron model data, but also actual biologically recorded *C. Elegans* neuron activity data. Comparing our method with that obtained by a reservoir computing method, we have achieved a higher count of TP (21 compared to 14) while keeping the same number of FPs (4). In summary, our results so far propose a newly more generalized method for connectivity estimation for neuron activity data, which is crucial for research in neuronal connectomics. For comparison, our best connectivity matrix with a threshold of 0.2 is shown in Fig. 10A, together with the GT connectivity in Fig. 10B. We show a receiver operating characteristics (ROC) curve in Fig. S24, where we systematically vary the binarization threshold. The result is that LCC outperforms the reservoir computing-based method for relatively large thresholds, leading to a lower false positive rate and a higher true positive rate. In Fig. S25 A, we show the Pearson correlation coefficient for different algorithms studied in this paper. It becomes apparent that all methods perform similarly, with relatively low values for the correlation. Similar results hold for the precision and the F1 score, but the values are higher overall, highlighting the need for adequate measures when considering binary GT matrices. (Fig. S25 B and C). The best performance for Pearson Correlation is shown by cut LCC, followed by correlation. This shows that reliably estimating *C. elegans* connectivity remains a challenge for our algorithms.

**Figure 10:**
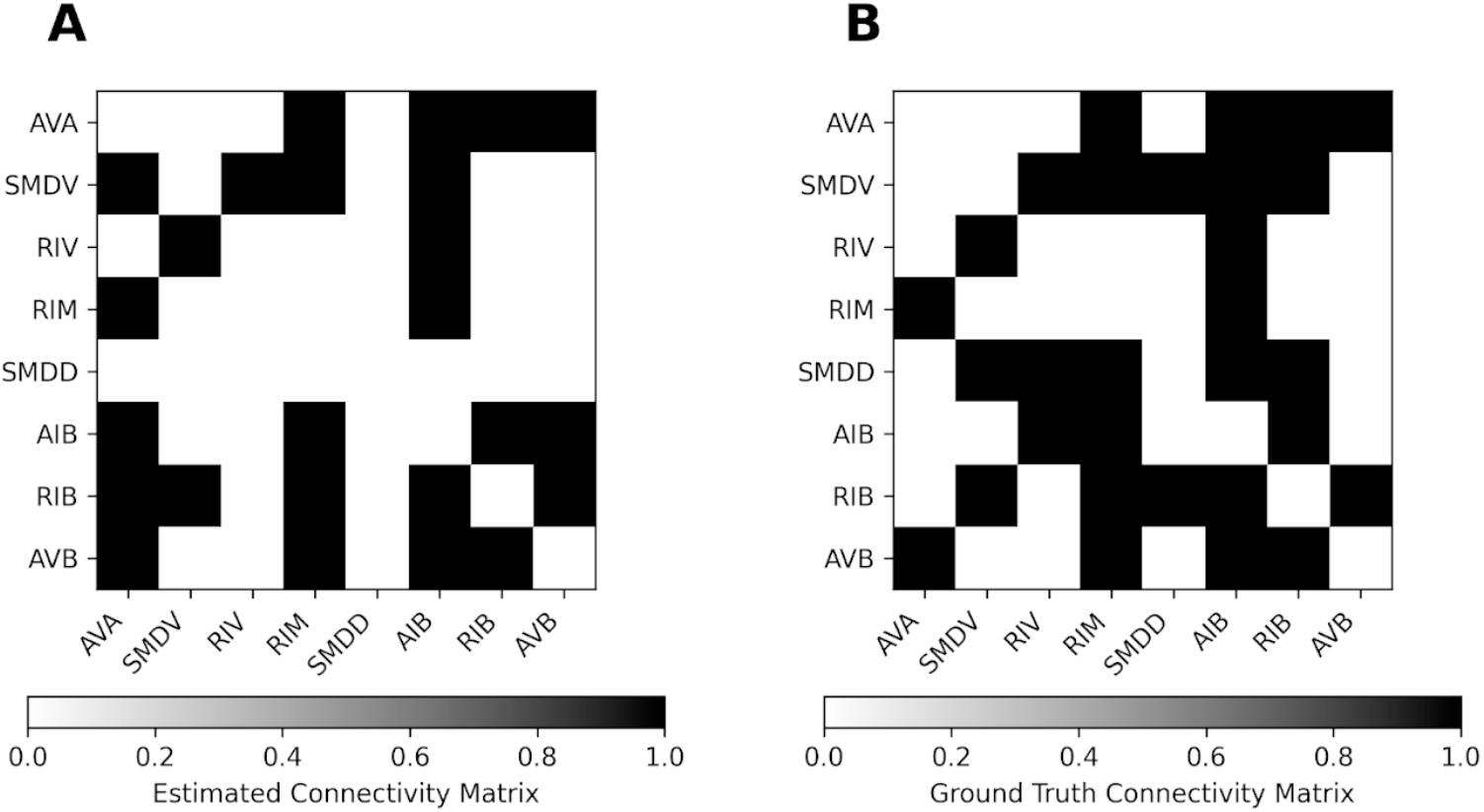
Comparison of the GT and EStC connectivity for the 8 most active motor neurons of *C. Elegans*. Panel A shows the EStC matrix using the LCC method with a binarization: values above a threshold of 0.2 are clipped to 1 and values below it to 0. Panel B shows the GT connectivity matrix for the 8 most active motor neurons of *C. elegans*. Comparing the matrices shows a TP count of 21 and an FP count of 4 as indicated in Fig. 9 by the black dashed line.

## 4 Discussion

The goal of this paper was twofold: first, we developed and validated a method to estimate the EC of the network from observed neural activity data, without relying on prior knowledge of the network topology or dynamics. Second, we systematically compared the performance of our LCC method with other methods from the broad classes of correlation-based (standard correlation and Partial Correlation), derivative-based (DDC) and information theory-based (NTE) measures in a non-linear neural mass model with delays and in a linear single-neuron model in the form of coupled Ornstein-Uhlenbeck processes. The first goal was pursued in two steps, by first inferring the underlying SC of the system and comparing it to GT data, and second, for the Hopf model, using the SC in conjunction with a model of the local node dynamics to recreate the observed activity patterns.

Our approach for estimating the SC of the system, LCC, is based on lagged correlation analysis, which first estimates the directionality and then the strength of connections between neurons by finding the optimal time lag that maximizes the cross-correlation between the neurons. Furthermore, this method was used in conjunction with the recently published method of DDC, which uses a derivative-based approach to infer EC. We then applied the different methods to synthetic data generated by a Hopf neuron model and a coupled linear dynamical system to validate the methods. The estimation algorithm was finally applied to biological data derived from the eight most active motor neurons of the nematode *C. Elegans*, a well-known model organism for connectomics with a known GT connectivity matrix.

Our results showed that our new method performed well in estimating the connectivity of the Hopf model, especially when the networks were sparsely connected. We further compared our method with other established methods such as pairwise correlation, and DDC. For sparse connectivity, our method outperformed others in terms of Pearson correlation between the EStC and GT connectivity matrices. For denser connectivity, when more bidirectional connections are present, standard correlation-based methods show comparable performance to our LCC-based methods. This is expected, because, in denser networks, estimating the directionality of pairwise connections is more difficult due to the influence of third nodes. Therefore, our LCC-based methods do not outperform standard correlation-based measures in this case. For the Hopf model, our LCC-based methods performed better than DDC-based methods. We systematically studied the dependence of the estimation performance on the system parameters, including the dynamical regime, noise parameters and the delay distribution.

In contrast to these findings for the Hopf model, LCC did not outperform DDC-based methods in a canonical simple neuron model (Eq. 8). The reason is given by the fact that DDC, being based on linear model equations without delay, is the least squares error estimator of the system given by Eq. 8, and hence, the performance of DDC-based methods should be high given a sufficient amount of data. Our LCC-based methods, however, still outperformed standard correlation-based measures, albeit not by a large margin (Figs. 5-7). For low connectivity, they also slightly outperformed NTE, but at a drastically lower computational cost (SI, Fig. S19).

We then demonstrated that our method could be used to recreate the system dynamics by using the estimated SC matrix as the new connectivity for a Hopf model with the same input parameters used for the generation of the GT data. We found that this produced high correlations between the original and regenerated data, indicating that our approach captured the essential dynamical features encoded in the connectivity matrix. Thus, our approach could be used to bridge the gap between SC and EC in this scenario.

Finally, we applied our method to *C. Elegans* activity data and found that it could infer the connectivity better than currently used methods such as the recently proposed reservoir computing approach by [48]. Our method achieved a higher count of true positives at the same amount of false positives compared to the reservoir computing approach. This finding suggests that our method is robust and reliable in estimating connectivity from biological data, which often struggles with noise and low temporal resolution.

Our work contributes to the field of computational connectomics by proposing a method to estimate SC, which can then be used in conjunction with a model of local node dynamics to derive the effective connectivity of the system and recreate the observed activity patterns. The two-step approach is surprisingly simple, which endows it with generality and flexibility, allowing for an application to different types of neural activity data. Another advantage of our LCC method is that we need no prior assumptions about network topology or node dynamics, in contrast to methods involving DDC, which assume linear or non-linear node dynamics of a specific type and no external input. As such, our LCC method, like established correlation-based methods for inferring connectivity, is parameter free. The thresholded LCC method needs one input parameter, namely the threshold, above which potential links are considered connections.

To give a practical guideline on how to proceed in a minimal scenario, where we only have access to GT neural activity data from a specific neuronal population of a given known size, we here outline critical steps for the selection of an estimation method (Fig. 11). In light of the results presented in this paper, different methods should be used for different dynamics and network sizes and sparsities. We always assume that our networks are small and/or sparse. We further assume that we have access to all relevant nodes (i.e. our observation of the network is complete) and that the sampled signal has a high temporal resolution. Next, with the help of auto- and cross-correlograms, the delay distribution in the network should be determined. For non-linear dynamics with small delays, we recommend the LCC method, possibly in conjunction with DDC. For linear dynamics with large or small/vanishing delays, we recommend the DDC method. For the cases not covered by these two (non-linear with large delays and linear with small delays), we recommend using the thresholded DDC method. If the network under consideration is not sparse, methods moving beyond pairwise interactions need to be considered. These methods are based on information theory (IT), for example, multivariate transfer entropy [65]. This method is also expected to work in larger and denser networks, albeit at a much higher computational cost.

**Figure 11:**
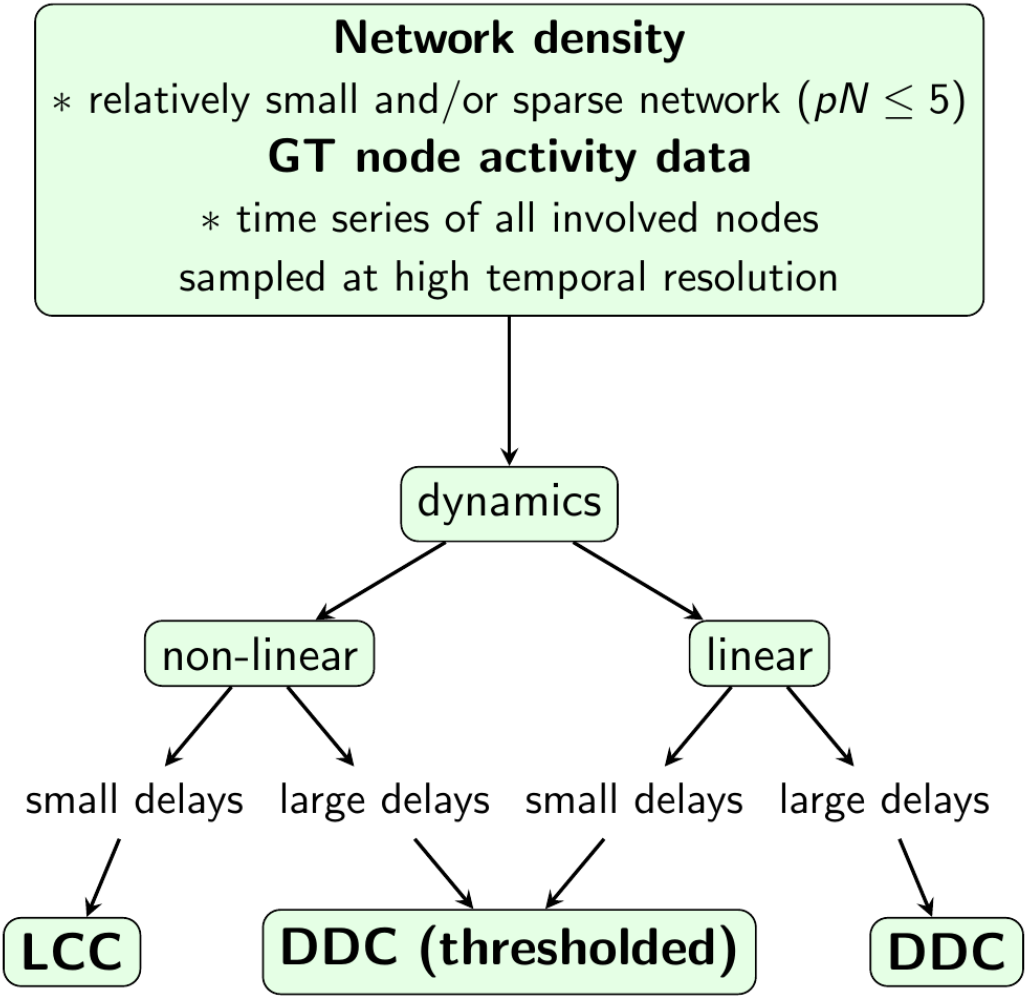
Guide for method selection to estimate connectivity from activity in neural networks. The input to all methods is recorded GT activity data, for which assumptions on the presence or absence of delays and underlying dynamical model need to be made. Also, an initial estimate about the network size and its density should be made. With these inputs and assumptions, we can proceed with a choice of estimation method. For non-linear dynamics and small delays, LCC is the preferred method, whereas DDC is the preferred method for linear dynamics with small/ absent or large delays. For non-linear dynamics with large delays, the thresholded DDC method is preferred.

How can these methods be improved? Clearly, dealing with the problem of partially observed dynamics remains a challenge. Even if our LCC method does not need the assumption of full observability, it certainly aids performance when network dynamics are generated intrinsically, and not driven from outside. When strong external driving is present, our approach can still result in errors arising from confounder motifs (see Introduction). To alleviate this problem, either the network needs to be observed more completely or external input must be taken into account in the formulation of the estimation algorithm, as done in [67]. Another direction for future research is to estimate connectivity in networks of inhibitory neurons [67]. Initial results show that using LCC in these networks results in EStC matrices that have a complementary structure compared to the GT matrix. This is not surprising, because two neurons that are strongly coupled by an inhibitory connection will show dynamics that is anti-correlated, and hence, we expect that a low entry in the EStC matrix actually reflects the presence of a strong inhibitory connection.

Another area for future work would be to combine our approach with methods from simulation-based inference [44], [70], [71]. Concretely, in the absence of GT connectivity, our LCC-based methods could be used to estimate structural connectivity. This connectivity could be fed into a mechanistic model of neural activity serving as a system generating the empirical activity data. Its parameters, but not the connectivity itself, could then be estimated using methods from simulation-based inference. When these parameters are found, the system identification loop is closed: the data can be recreated by forward-simulating the estimated model with the estimated SC matrix. Alternatively, for small networks, dynamical causal modeling (DCM) [72] could be used. In case of good agreement of the simulated, recreated traces with the original activity data, this step closes the loop and bridges the gap between SC and EC. In other words, a system that generates the data has been successfully identified. These exciting questions are left for future work.

## Supporting information

SI with 25 supplemental figures

## 6 Author Contributions

Conceptualization NL, WB, LK, JB, CCH

Data curation NL

Formal analysis NL, WB

Funding acquisition CCH

Investigation NL, WB, CCH, LK

Methodology NL, WB, LK

Project administration WB, CCH

Resources NL, CCH

Software NL

Supervision WB, CCH

Validation NL, WB

Visualization NL

Writing – original draft NL, WB

Writing – review & editing NL, WB, LK, JB, CCH

## 8 Acknowledgments

The authors would like to thank Kayson Fakhar, Alexander Schaum, Fatemeh Hadaeghi, Arnaud Messé, Gorka Zamora-López and Heike Siebert for useful comments.

## 9 Data Availability Statement

Code used in this study can be found under www.github.com/UKE-ICNS/LCC. The datasets generated or analyzed in the current study are available from the corresponding author on reasonable request.

## Bibliography

[1] F. V. Farahani, W. Karwowski, and N. R. Lighthall, “Application of Graph Theory for Identifying Connectivity Patterns in Human Brain Networks: A Systematic Review,” Front. Neurosci., vol. 13, 2019, Accessed: Jan. 24, 2024. [Online]. Available: https://www.frontiersin.org/articles/10.3389/fnins.2019.00585

[2] E. Bullmore and O. Sporns, “Complex brain networks: graph theoretical analysis of structural and functional systems,” Nat. Rev. Neurosci., vol. 10, no. 3, Art. no. 3, Mar. 2009, doi: 10.1038/nrn2575.

[3] C.-H. Yeh, D. K. Jones, X. Liang, M. Descoteaux, and A. Connelly, “Mapping Structural Connectivity Using Diffusion MRI: Challenges and Opportunities,” J. Magn. Reson. Imaging, vol. 53, no. 6, pp. 1666–1682, 2021, doi: 10.1002/jmri.27188.

[4] K. H. Maier-Hein et al., “The challenge of mapping the human connectome based on diffusion tractography,” Nat. Commun., vol. 8, no. 1, Art. no. 1, Nov. 2017, doi: 10.1038/s41467-017-01285-x.

[5] O. Sporns, G. Tononi, and R. Kötter, “The Human Connectome: A Structural Description of the Human Brain,” PLOS Comput. Biol., vol. 1, no. 4, p. e42, Sep. 2005, doi: 10.1371/journal.pcbi.0010042.

[6] J. G. White, E. Southgate, J. N. Thomson, and S. Brenner, “The structure of the nervous system of the nematode Caenorhabditis elegans,” Philos. Trans. R. Soc. Lond. B. Biol. Sci., vol. 314, no. 1165, pp. 1–340, Nov. 1986, doi: 10.1098/rstb.1986.0056.

[7] G. Yan et al., “Network control principles predict neuron function in the Caenorhabditis elegans connectome,” Nature, vol. 550, no. 7677, Art. no. 7677, Oct. 2017, doi: 10.1038/nature24056.

[8] M. Winding et al., “The connectome of an insect brain,” Science, vol. 379, no. 6636, p. eadd9330, Mar. 2023, doi: 10.1126/science.add9330.

[9] K. J. Friston, C. D. Frith, P.F. Liddle, and R.S.J. Frackowiak, “Functional Connectivity: The Principal-Component Analysis of Large (PET) Data Sets,” J. Cereb. Blood Flow Metab., vol. 13, no. 1, pp. 5–14, Jan. 1993, doi: 10.1038/jcbfm.1993.4.

[10] I. H. Stevenson, J. M. Rebesco, L. E. Miller, and K. P. Körding, “Inferring functional connections between neurons,” Curr. Opin. Neurobiol., vol. 18, no. 6, pp. 582–588, Dec. 2008, doi: 10.1016/j.conb.2008.11.005.

[11] K. J. Friston, “Functional and effective connectivity in neuroimaging: A synthesis,” Hum. Brain Mapp., vol. 2, no. 1–2, pp. 56–78, 1994, doi: 10.1002/hbm.460020107.

[12] K. J. Friston, “Functional and effective connectivity: a review,” Brain Connect., vol. 1, no. 1, pp. 13–36, 2011, doi: 10.1089/brain.2011.0008.

[13] P. A. Valdes-Sosa, A. Roebroeck, J. Daunizeau, and K. Friston, “Effective connectivity: Influence, causality and biophysical modeling,” NeuroImage, vol. 58, no. 2, pp. 339–361, Sep. 2011, doi: 10.1016/j.neuroimage.2011.03.058.

[14] P. C. Klink et al., “Combining brain perturbation and neuroimaging in non-human primates,” NeuroImage, vol. 235, p. 118017, Jul. 2021, doi: 10.1016/j.neuroimage.2021.118017.

[15] K. Fakhar and C. C. Hilgetag, “Systematic perturbation of an artificial neural network: A step towards quantifying causal contributions in the brain,” PLOS Comput. Biol., vol. 18, no. 6, p. e1010250, Jun. 2022, doi: 10.1371/journal.pcbi.1010250.

[16] M. Gilsonet et al., “Model-based whole-brain effective connectivity to study distributed cognition in health and disease,” Netw. Neurosci., vol. 4, no. 2, pp. 338–373, Apr. 2020, doi: 10.1162/netn_a_00117.

[17] M. Gilson, R. Moreno-Bote, A. Ponce-Alvarez, P. Ritter, and G. Deco, “Estimation of Directed Effective Connectivity from fMRI Functional Connectivity Hints at Asymmetries of Cortical Connectome,” PLOS Comput. Biol., vol. 12, no. 3, p. e1004762, Mar. 2016, doi: 10.1371/journal.pcbi.1004762.

[18] G. Nolte et al, “Robustly Estimating the Flow Direction of Information in Complex Physical Systems,” Phys. Rev. Lett., vol. 100, no. 23, p. 234101, Jun. 2008, doi: 10.1103/PhysRevLett.100.234101.

[19] C. J. Stam, G. Nolte, and A. Daffertshofer, “Phase lag index: Assessment of functional connectivity from multi channel EEG and MEG with diminished bias from common sources,” Hum. Brain Mapp., vol. 28, no. 11, pp. 1178–1193, 2007, doi: 10.1002/hbm.20346.

[20] A. Fornito, A. Zalesky, and E. T. Bullmore, Eds., “Front Matter,” in Fundamentals of Brain Network Analysis, San Diego: Academic Press, 2016, pp. i–ii. doi: 10.1016/B978-0-12-407908-3.09996-9.

[21] M. Rubinov and O. Sporns, “Complex network measures of brain connectivity: Uses and interpretations,” NeuroImage, vol. 52, no. 3, pp. 1059–1069, Sep. 2010, doi: 10.1016/j.neuroimage.2009.10.003.

[22] Y. Chen, B. Q. Rosen, and T. J. Sejnowski, “Dynamical differential covariance recovers directional network structure in multiscale neural systems,” Proc. Natl. Acad. Sci., vol. 119, no. 24, p. e2117234119, Jun. 2022, doi: 10.1073/pnas.2117234119.

[23] K. Fakhar, F. Hadaeghi, and C. C. Hilgetag, “Causal Influences Decouple From Their Underlying Network Structure In Echo State Networks,” in2022 International Joint Conference on Neural Networks (IJCNN), Jul. 2022, pp. 1–8. doi: 10.1109/IJCNN55064.2022.9892782.

[24] C. J. Honey, R. Kötter, M. Breakspear, and O. Sporns, “Network structure of cerebral cortex shapes functional connectivity on multiple time scales,” Proc. Natl. Acad. Sci., vol. 104, no. 24, pp. 10240–10245, Jun. 2007, doi: 10.1073/pnas.0701519104.

[25] H. C. Tam, E. S. C. Ching, and P.-Y. Lai, “Reconstructing networks from dynamics with correlated noise,” Phys. Stat. Mech. Its Appl., vol. 502, pp. 106–122, Jul. 2018, doi: 10.1016/j.physa.2018.02.166.

[26] L. Faes and G. Nollo, “Extended causal modeling to assess Partial Directed Coherence in multiple time series with significant instantaneous interactions,” Biol. Cybern., vol. 103, no. 5, pp. 387–400, Nov. 2010, doi: 10.1007/s00422-010-0406-6.

[27] D. Poli, V. P. Pastore, S. Martinoia, and P. Massobrio, “From functional to structural connectivity using partial correlation in neuronal assemblies,” J. Neural Eng., vol. 13, no. 2, p. 026023, Feb. 2016, doi: 10.1088/1741-2560/13/2/026023.

[28] M. Eichler, R. Dahlhaus, and J. Sandkühler, “Partial correlation analysis for the identification of synaptic connections,” Biol. Cybern., vol. 89, no. 4, pp. 289–302, Oct. 2003, doi: 10.1007/s00422-003-0400-3.

[29] F. Randi, A. K. Sharma, S. Dvali, and A. M. Leifer, “Neural signal propagation atlas of Caenorhabditis elegans,” Nature, vol. 623, no. 7986, Art. no. 7986, Nov. 2023, doi: 10.1038/s41586-023-06683-4.

[30] E. Marder, “Neuromodulation of Neuronal Circuits: Back to the Future,” Neuron, vol. 76, no. 1, pp. 1–11, Oct. 2012, doi: 10.1016/j.neuron.2012.09.010.

[31] M. Timme and J. Casadiego, “Revealing networks from dynamics: an introduction,” J. Phys. Math. Theor., vol. 47, no. 34, p. 343001, Aug. 2014, doi: 10.1088/1751-8113/47/34/343001.

[32] P. Hagmannet et al., “Mapping the Structural Core of Human Cerebral Cortex,” PLOS Biol., vol. 6, no. 7, p. e159, Jul. 2008, doi: 10.1371/journal.pbio.0060159.

[33] C. Lainscsek, C. E. Gonzalez, A. L. Sampson, S. S. Cash, and T. J. Sejnowski, “Causality detection in cortical seizure dynamics using cross-dynamical delay differential analysis,” Chaos Interdiscip. J. Nonlinear Sci., vol. 29, no. 10, p. 101103, Oct. 2019, doi: 10.1063/1.5126125.

[34] G. C. Goodwin and R. L. Payne, Dynamic System Identification: Experiment Design and Data Analysis. Academic Press, 1977.

[35] S. G. Shandilya and M. Timme, “Inferring network topology from complex dynamics,” New J. Phys., vol. 13, no. 1, p. 013004, Jan. 2011, doi: 10.1088/1367-2630/13/1/013004.

[36] J. K. Lappalainenet et al., “Connectome-constrained networks predict neural activity across the fly visual system,” Nature, pp. 1–9, Sep. 2024, doi: 10.1038/s41586-024-07939-3.

[37] A. A. Prinz, D. Bucher, and E. Marder, “Similar network activity from disparate circuit parameters,” Nat. Neurosci., vol. 7, no. 12, Art. no. 12, Dec. 2004, doi: 10.1038/nn1352.

[38] M. Deistler, J. H. Macke, and P. J. Gonçalves, “Energy-efficient network activity from disparate circuit parameters,” Proc. Natl. Acad. Sci., vol. 119, no. 44, p. e2207632119, Nov. 2022, doi: 10.1073/pnas.2207632119.

[39] B. Weissbourd, T. Momose, A. Nair, A. Kennedy, B. Hunt, and D. J. Anderson, “A genetically tractable jellyfish model for systems and evolutionary neuroscience,” Cell, vol. 184, no. 24, pp. 5854–5868.e20, Nov. 2021, doi: 10.1016/j.cell.2021.10.021.

[40] F. Pallasdies, S. Goedeke, W. Braun, and R.-M. Memmesheimer, “From single neurons to behavior in the jellyfish Aurelia aurita,” eLife, vol. 8, p. e50084, Dec. 2019, doi: 10.7554/eLife.50084.

[41] C. Dupre and R. Yuste, “Non-overlapping Neural Networks in Hydra vulgaris,” Curr. Biol., vol. 27, no. 8, pp. 1085–1097, Apr. 2017, doi: 10.1016/j.cub.2017.02.049.

[42] J. Bielecki, S. K. Dam Nielsen, G. Nachman, and A. Garm, “Associative learning in the box jellyfish Tripedalia cystophora,” Curr. Biol., vol. 33, no. 19, pp. 4150–4159.e5, Oct. 2023, doi: 10.1016/j.cub.2023.08.056.

[43] J. Ladenbauer, S. McKenzie, D. F. English, O. Hagens, and S. Ostojic, “Inferring and validating mechanistic models of neural microcircuits based on spike-train data,” Nat. Commun., vol. 10, no. 1, Art. no. 1, Oct. 2019, doi: 10.1038/s41467-019-12572-0.

[44] P. J. Gonçalves et al., “Training deep neural density estimators to identify mechanistic models of neural dynamics,” eLife, vol. 9, p. e56261, Sep. 2020, doi: 10.7554/eLife.56261.

[45] D. S. Bassett and O. Sporns, “Network neuroscience,” Nat. Neurosci., vol. 20, no. 3, Art. no. 3, Mar. 2017, doi: 10.1038/nn.4502.

[46] G. Deco, M. L. Kringelbach, V. K. Jirsa, and P. Ritter, “The dynamics of resting fluctuations in the brain: metastability and its dynamical cortical core,” Sci. Rep., vol. 7, no. 1, Art. no. 1, Jun. 2017, doi: 10.1038/s41598-017-03073-5.

[47] C. Cakan and K. Obermayer, “Biophysically grounded mean-field models of neural populations under electrical stimulation,” PLOS Comput. Biol., vol. 16, no. 4, p. e1007822, Apr. 2020, doi: 10.1371/journal.pcbi.1007822.

[48] A. Banerjee, S. Chandra, and E. Ott, “Network inference from short, noisy, low time-resolution, partial measurements: Application to C. elegans neuronal calcium dynamics,” Proc. Natl. Acad. Sci., vol. 120, no. 12, p. e2216030120, Mar. 2023, doi: 10.1073/pnas.2216030120.

[49] S. L. Brunton, J. L. Proctor, and J. N. Kutz, “Discovering governing equations from data by sparse identification of nonlinear dynamical systems,” Proc. Natl. Acad. Sci., vol. 113, no. 15, pp. 3932–3937, Apr. 2016, doi: 10.1073/pnas.1517384113.

[50] G. A. Cecchi, A. R. Rao, M. V. Centeno, M. Baliki, A. V. Apkarian, and D. R. Chialvo, “Identifying directed links in large scale functional networks: application to brain fMRI,” BMC Cell Biol., vol. 8, no. 1, p. S5, Jul. 2007, doi: 10.1186/1471-2121-8-S1-S5.

[51] J. Schieferet et al., “From correlation to causation: Estimating effective connectivity from zero-lag covariances of brain signals,” PLOS Comput. Biol., vol. 14, no. 3, p. e1006056, Mar. 2018, doi: 10.1371/journal.pcbi.1006056.

[52] Z. K. Tian, K. Chen, S. Li, D. W. McLaughlin, and D. Zhou, “Causal connectivity measures for pulse-output network reconstruction: Analysis and applications,” Proc. Natl. Acad. Sci., vol. 121, no. 14, p. e2305297121, Apr. 2024, doi: 10.1073/pnas.2305297121.

[53] A. Mitra, A. Z. Snyder, C. D. Hacker, and M. E. Raichle, “Lag structure in resting-state fMRI,” J. Neurophysiol., vol. 111, no. 11, pp. 2374–2391, Jun. 2014, doi: 10.1152/jn.00804.2013.

[54] A. Mitra, A. Z. Snyder, T. Blazey, and M. E. Raichle, “Lag threads organize the brain’s intrinsic activity,” Proc. Natl. Acad. Sci., vol. 112, no. 17, pp. E2235–E2244, Apr. 2015, doi: 10.1073/pnas.1503960112.

[55] B. Park, W. M. Shim, O. James, and H. Park, “Possible links between the lag structure in visual cortex and visual streams using fMRI,” Sci. Rep., vol. 9, no. 1, p. 4283, Mar. 2019, doi: 10.1038/s41598-019-40728-x.

[56] J. Heyse, L. Sheybani, S. Vulliémoz, and P. van Mierlo, “Evaluation of Directed Causality Measures and Lag Estimations in Multivariate Time-Series,” Front. Syst. Neurosci., vol. 15, p. 620338, Jan. 2021, doi: 10.3389/fnsys.2021.620338.

[57] T. Schreiber, “Measuring Information Transfer,” Phys. Rev. Lett., vol. 85, no. 2, pp. 461–464, Jul. 2000, doi: 10.1103/PhysRevLett.85.461.

[58] S. McCabeet et al., “netrd: A library for network reconstruction and graph distances,” J. Open Source Softw., vol. 6, no. 62, p. 2990, Jun. 2021, doi: 10.21105/joss.02990.

[59] C. Cakan, N. Jajcay, and K. Obermayer, “neurolib: A Simulation Framework for Whole-Brain Neural Mass Modeling,” Cogn. Comput., vol. 15, no. 4, pp. 1132–1152, Jul. 2023, doi: 10.1007/s12559-021-09931-9.

[60] S. Coombes, P. Beim Graben, and R. Potthast, “Tutorial on Neural Field Theory,” inNeural Fields: Theory and Applications, S. Coombes, P. Beim Graben, R. Potthast, and J. Wright, Eds., Berlin, Heidelberg: Springer, 2014, pp. 1–43. doi: 10.1007/978-3-642-54593-1_1.

[61] A. Ponce-Alvarez and G. Deco, “The Hopf whole-brain model and its linear approximation,” Sci. Rep., vol. 14, no. 1, p. 2615, Jan. 2024, doi: 10.1038/s41598-024-53105-0.

[62] F. Svaraet et al., “Automated synapse-level reconstruction of neural circuits in the larval zebrafish brain,” Nat. Methods, vol. 19, no. 11, Art. no. 11, Nov. 2022, doi: 10.1038/s41592-022-01621-0.

[63] J. Sun, D. Taylor, and E. M. Bollt, “Causal Network Inference by Optimal Causation Entropy,” SIAM J. Appl. Dyn. Syst., vol. 14, no. 1, pp. 73–106, Jan. 2015, doi: 10.1137/140956166.

[64] L. Novelli and J. T. Lizier, “Inferring network properties from time series using transfer entropy and mutual information: Validation of multivariate versus bivariate approaches,” Netw. Neurosci., vol. 5, no. 2, pp. 373–404, May 2021, doi: 10.1162/netn_a_00178.

[65] L. Novelli, p. Wollstadt, P. Mediano, M. Wibral, and J. T. Lizier, “Large-scale directed network inference with multivariate transfer entropy and hierarchical statistical testing,” Netw. Neurosci., vol. 3, no. 3, pp. 827–847, Jul. 2019, doi: 10.1162/netn_a_00092.

[66] R. Kobayashiet et al., “Reconstructing neuronal circuitry from parallel spike trains,” Nat. Commun., vol. 10, no. 1, p. 4468, Oct. 2019, doi: 10.1038/s41467-019-12225-2.

[67] T. Nieus, D. Borgonovo, S. Diwakar, G. Aletti, and G. Naldi, “A multi-class logistic regression algorithm to reliably infer network connectivity from cell membrane potentials,” Front. Appl. Math. Stat., vol. 8, 2022, Accessed: Feb. 22, 2024. [Online]. Available: https://www.frontiersin.org/articles/10.3389/fams.2022.1023310

[68] V. Pernice and S. Rotter, “Reconstruction of sparse connectivity in neural networks from spike train covariances,” J. Stat. Mech. Theory Exp., vol. 2013, no. 03, p. P03008, Mar. 2013, doi: 10.1088/1742-5468/2013/03/P03008.

[69] S. Katoet et al., “Global Brain Dynamics Embed the Motor Command Sequence of Caenorhabditis elegans,” Cell, vol. 163, no. 3, pp. 656–669, Oct. 2015, doi: 10.1016/j.cell.2015.09.034.

[70] K. Cranmer, J. Brehmer, and G. Louppe, “The frontier of simulation-based inference,” Proc. Natl. Acad. Sci., vol. 117, no. 48, pp. 30055–30062, Dec. 2020, doi: 10.1073/pnas.1912789117.

[71] J. Boelts et al., “Simulation-based inference for efficient identification of generative models in computational connectomics,” PLOS Comput. Biol., vol. 19, no. 9, p. e1011406, Sep. 2023, doi: 10.1371/journal.pcbi.1011406.

[72] K. J. Friston, L. Harrison, and W. Penny, “Dynamic causal modelling,” NeuroImage, vol. 19, no. 4, pp. 1273–1302, Aug. 2003, doi: 10.1016/S1053-8119(03)00202-7.

