## Supplementary material for "Estimating effective connectivity in neural networks: comparison of derivative-based and correlation-based methods": SI with 25 supplemental figures

**for**

Note: if not mentioned otherwise, M=10 realizations (instead of M=100 in the main manuscript) per parameter value and method were used for the following figures where appropriate.

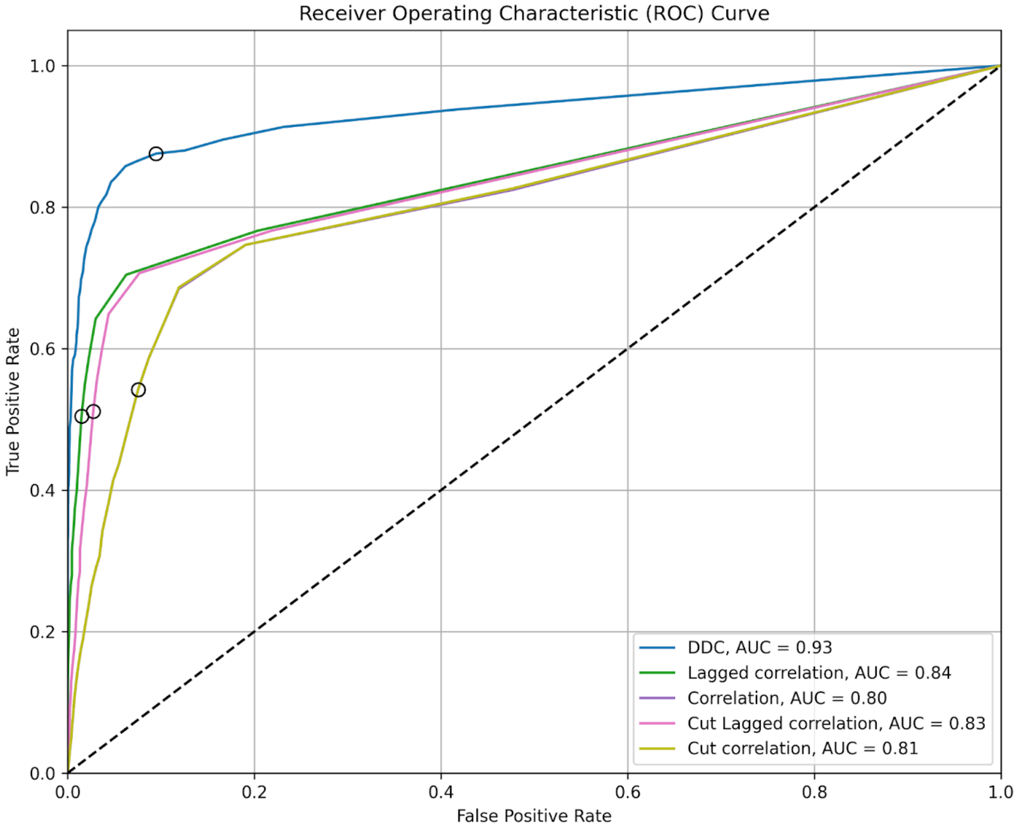


**Figure S1: Receiver-operator characteristics (ROC) curve for the different models using a Hopf network (N=10, p=0.1) and a fixed delay of 100 ms.** The parameter for the ROC curve is the binarization threshold for the estimated matrix, which increases from top right to bottom left. As seen in the ROC curve, DDC performed best for the large delays, with Lagged Correlation and cut Lagged Correlation performing better than standard correlations (AUC, area under the curve). The black circles indicate the binarization threshold of 0.1. The curves for ‘Correlation’ and ‘Cut correlation’ show substantial overlap.

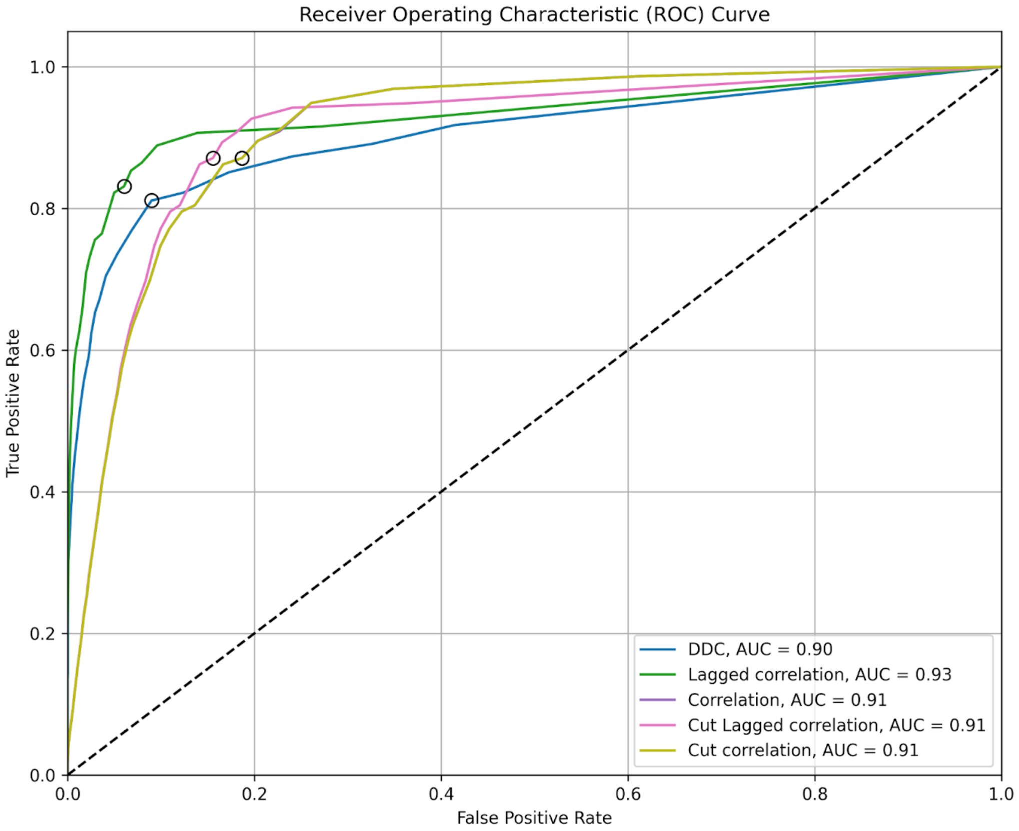


**Figure S2: Receiver-operator characteristics (ROC) curve for the different models using a Hopf network (N=10, p=0.1) and a fixed delay of 10 ms.** The parameter for the ROC curve was the binarization threshold for the estimated matrix. As seen in the ROC curve, Lagged correlation performed best overall with the largest area under the curve. Lagged correlation plus DDC, Lagged correlation thresholded and DDC thresholded were removed from the ROC calculations as they intrinsically already use a threshold. The black circles indicate the binarization threshold of 0.1. The curves for 'Correlation' and 'Cut correlation' show substantial overlap.

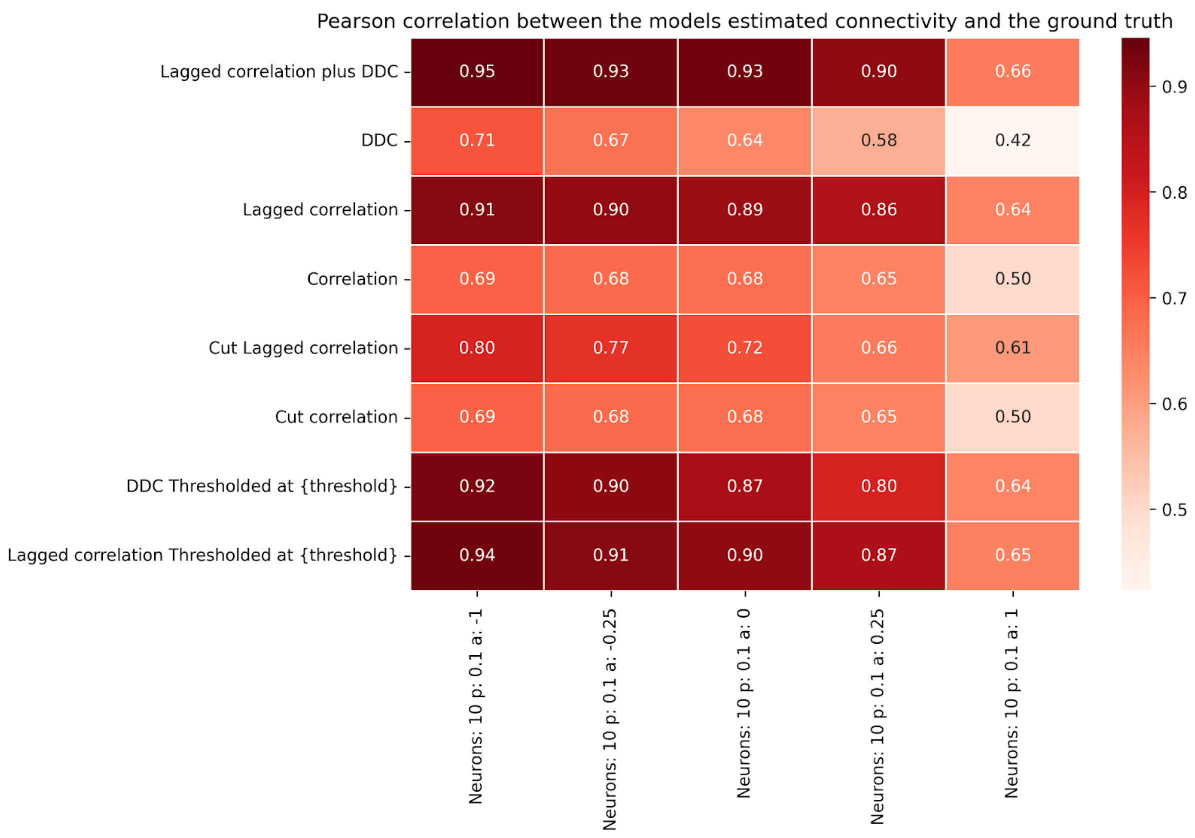


**Figure S3: Dependence of connectivity estimation performance in Hopf networks (N=10, p=0.1) on bifurcation parameter a for small delays.** Pearson correlation of connectivity estimation methods for a Hopf oscillator network (N = 10, p = 0.1, uniformly distributed delays [0 ms, 10 ms], as in Fig. 2 in the main manuscript) across bifurcation parameter a. Lagged correlation methods outperform thresholded DDC for a < 0. At a = 0.25, Lagged Correlation thresholded and Lagged Correlation plus DDC show better performance than DDC thresholded.

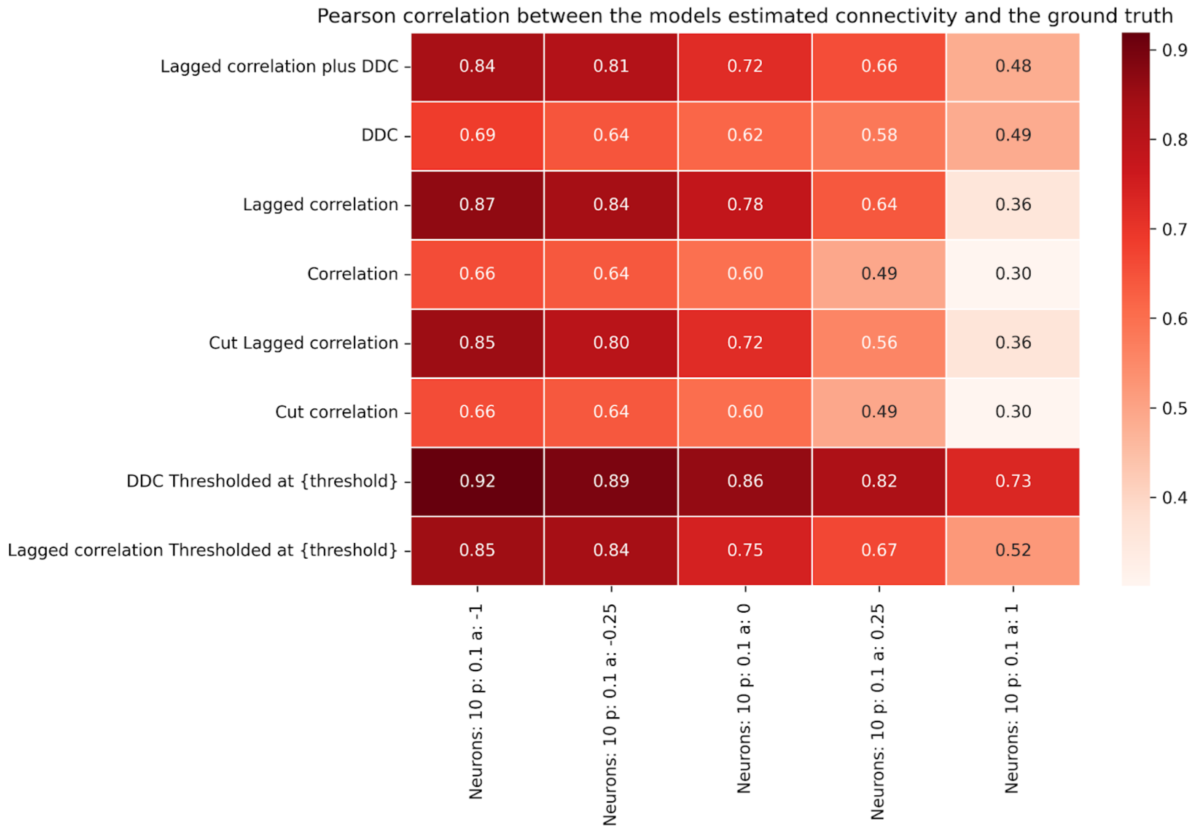


**Figure S4: Dependence of connectivity estimation performance in Hopf networks (N=10, p=0.1) on bifurcation parameter a for large delays.** Pearson correlation of different connectivity estimation methods for a Hopf oscillator network (N = 10, p = 0.1, uniformly distributed delays [0 ms, 100 ms]) across bifurcation parameter a ∈ [-1, 1]. Thresholded DDC shows best overall performance but declines with increasing a. Lagged and Cut Lagged correlations perform well for a ≤ 0 but deteriorate for a > 0, likely due to delays exceeding oscillation periods.

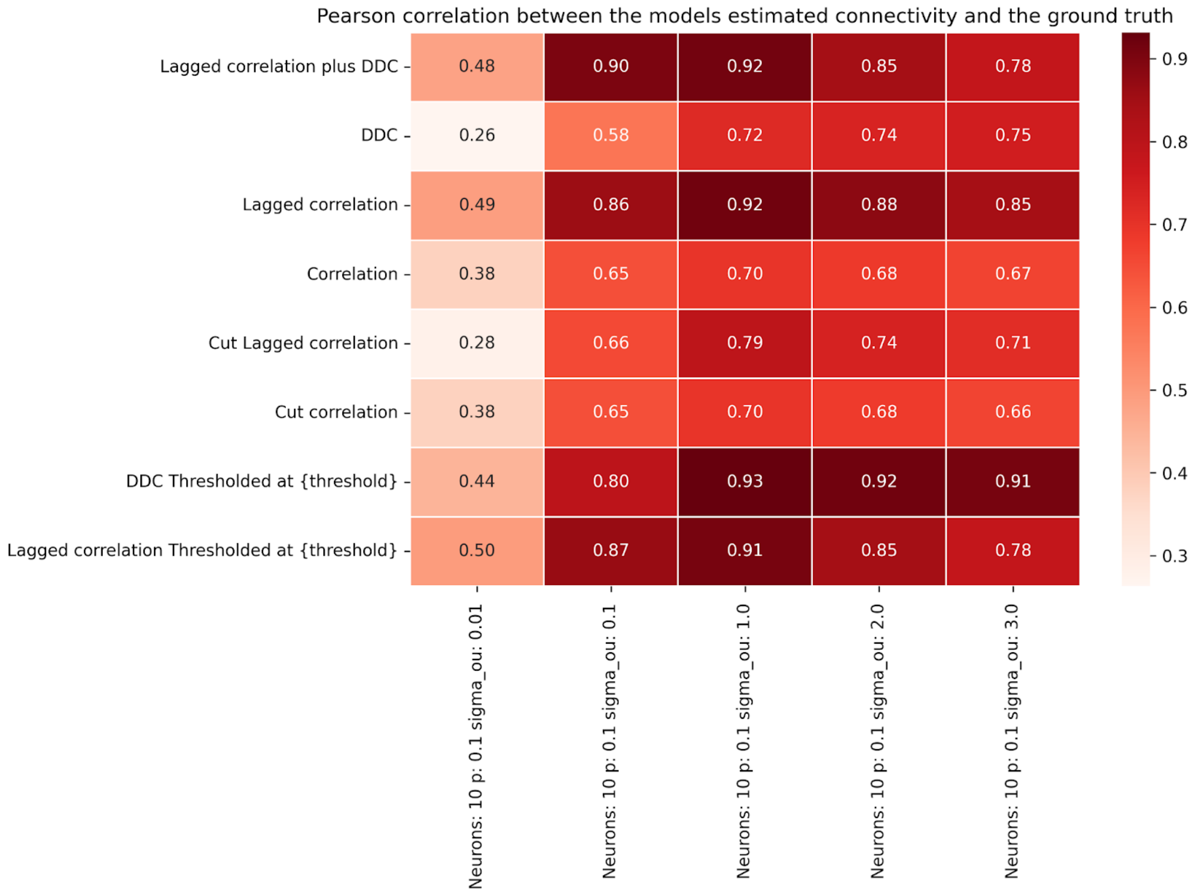


**Figure S5: Impact of Ornstein-Uhlenbeck noise strength on connectivity estimation performance in Hopf networks (N=10, p=0.1).** Parameters as in Fig. 2 of main manuscript, except for fixed delays (fixed at 10 ms) and different values for sigma_OU. Pearson correlation of connectivity estimation methods for a network (N = 10, p = 0.1) with varying Ornstein-Uhlenbeck noise (sigma_ou). Performance generally improves with increasing sigma_ou. At sigma_OU= 1, DDC thresholded outperforms lagged correlation methods. The default value for the main manuscript is sigma_OU = 0.1. At larger values for sigma_OU (approximately after sigma_OU = 1), the performance decreases. When sigma_OU reaches 5, the simulation does not produce interpretable results and connectivity estimation is not possible.

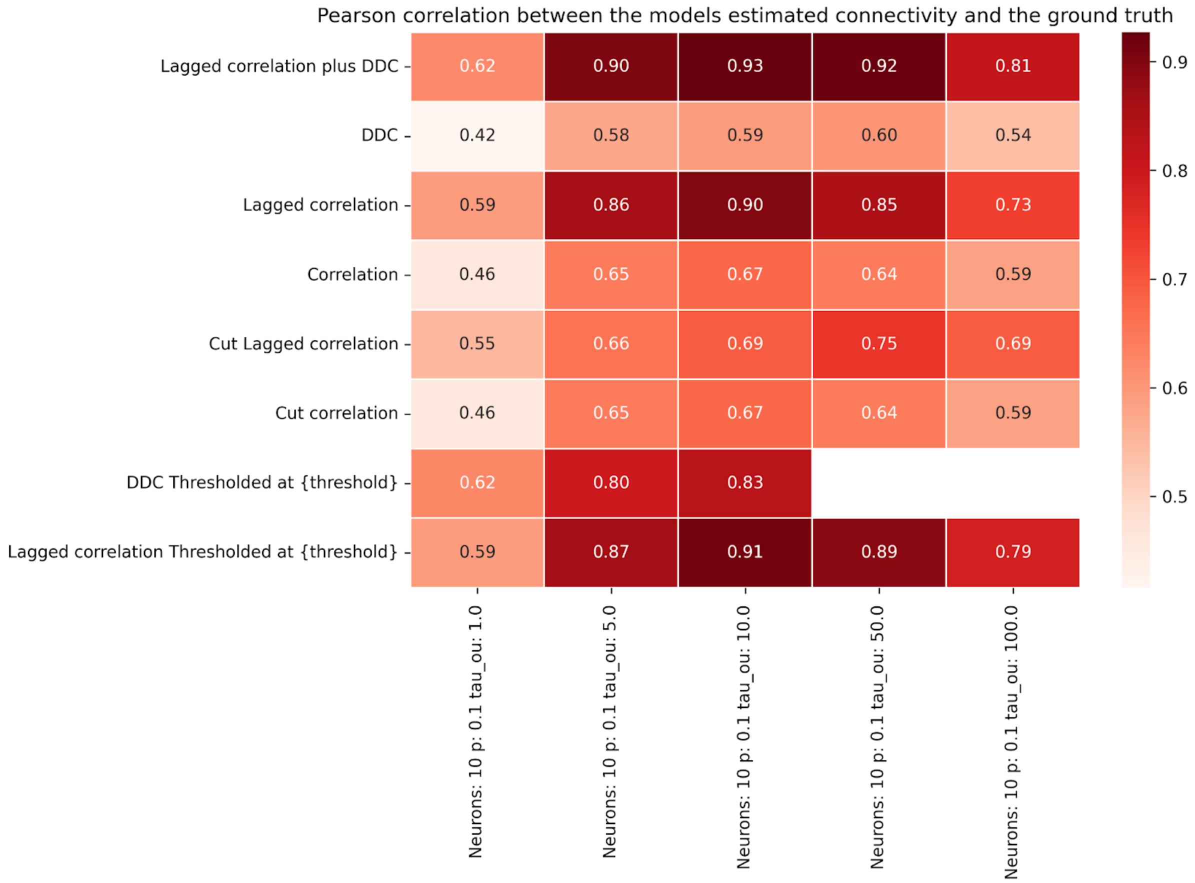


**Figure S6: Impact of Ornstein-Uhlenbeck time constant on connectivity estimation performance in Hopf networks (N=10, p=0.1).** Parameters as in Fig. 2 of main manuscript, except for fixed delays (fixed at 10 ms) and different values for tau_OU. Pearson correlation of connectivity estimation methods for a network (nN= 10, p = 0.1) with varying Ornstein-Uhlenbeck time constant (tau_ou). Similar to the results for sigma_OU, performance improves with increasing tau_OU, and then decreases (approximately after tau_OU = 10ms). For larger tau_OU, lagged correlation methods outperform others. Paper uses tau_OU= 5.0 ms. The white squares represent NaN values in the estimated matrix.

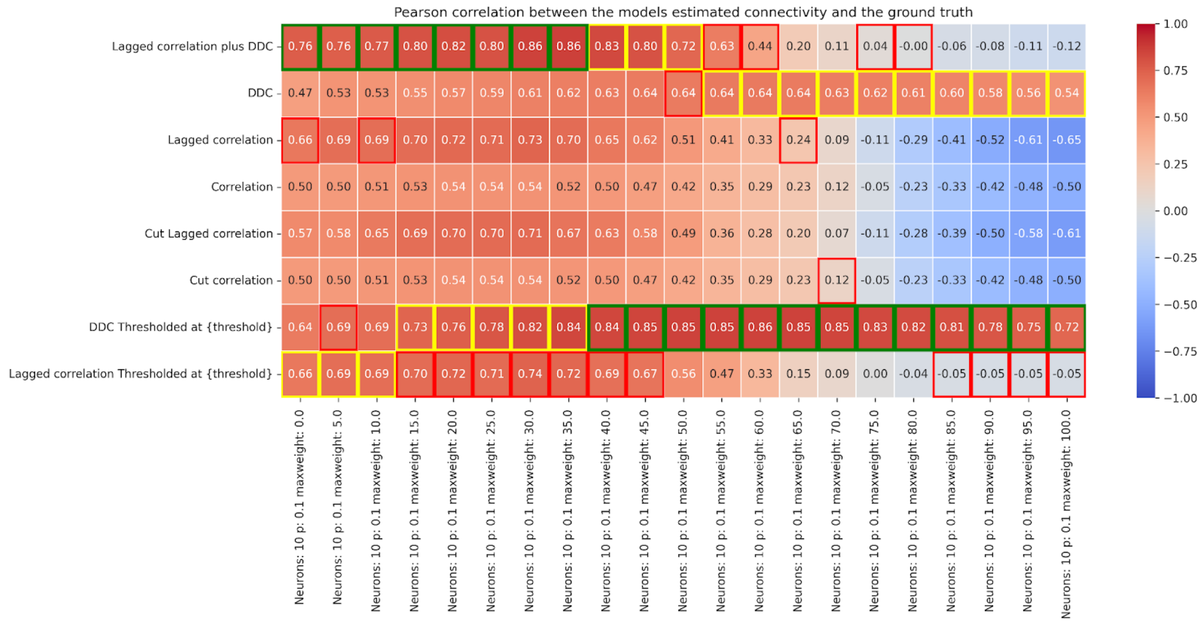


**Figure S7: Hopf model: Connectivity estimation performance of methods in relation to changing delays and a = 1.** The delay parameter increases from left to right. The value of maxweight indicates the fixed delay value in ms. Remaining parameter values, except delay and value for a, as in Fig. 2 in main manuscript. Highlighted in green is the best performing method for that specific delay. Yellow for the second best performing method and red for the third best.

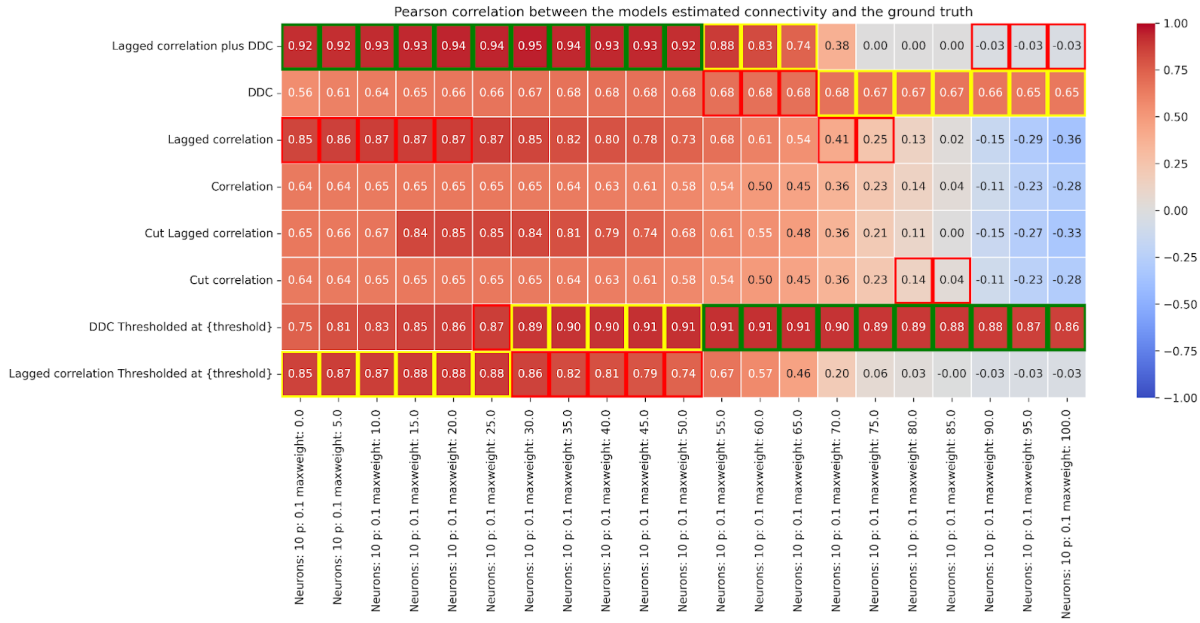


**Figure S8: Hopf model: Connectivity estimation performance of methods in relation to changing delays and a = 0.25.** Figure layout as in SI Fig. 13.

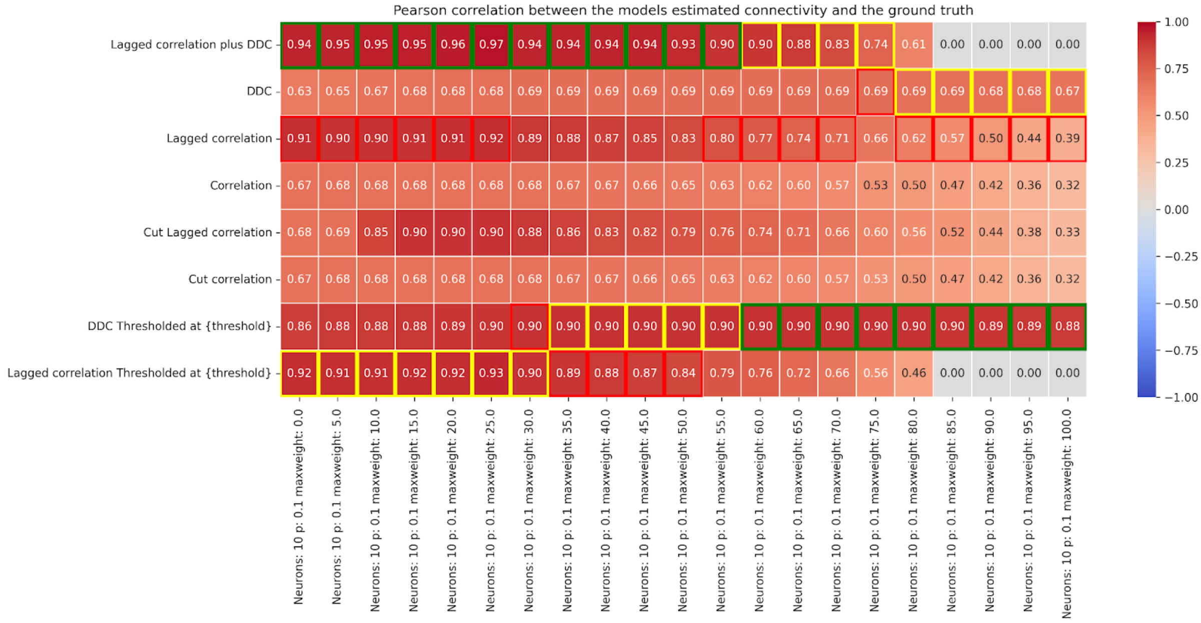


**Figure S9: Hopf model: Connectivity estimation performance of methods in relation to changing delays and a = 0.** Figure layout as in SI Fig. 13.

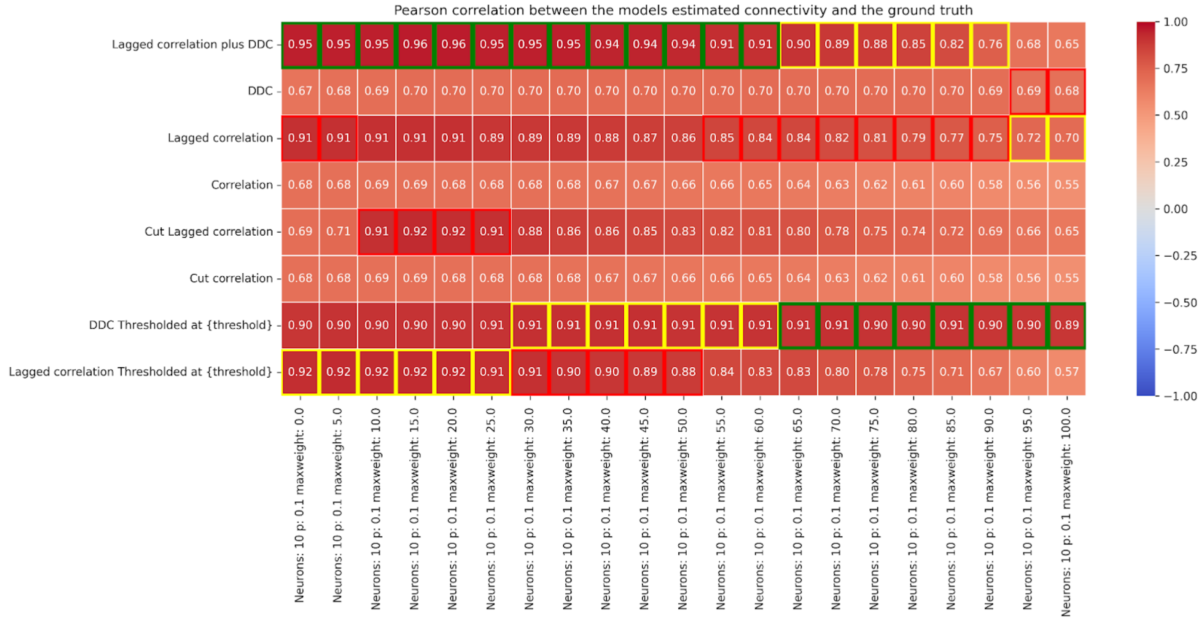


**Figure S10: Hopf model: Connectivity estimation performance of methods in relation to changing delays and a = - 0.25.** Figure layout as in SI Fig. 13.

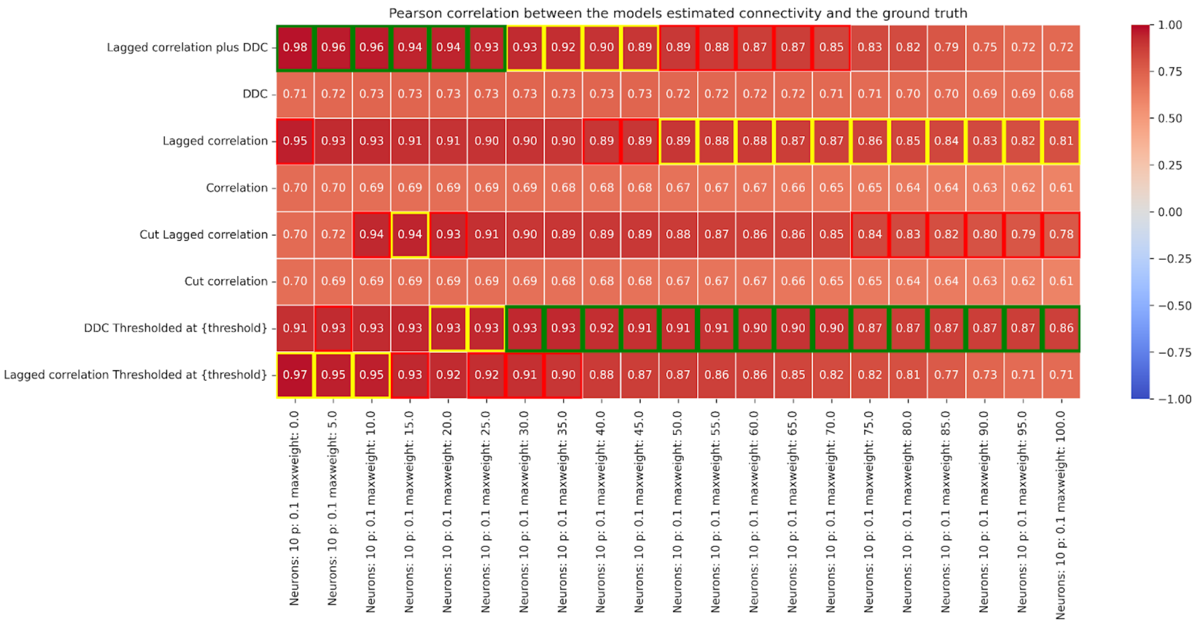


**Figure S11: Hopf model: Connectivity estimation performance of methods in relation to changing delays and a = - 1.** Figure layout as in SI Fig. 13.

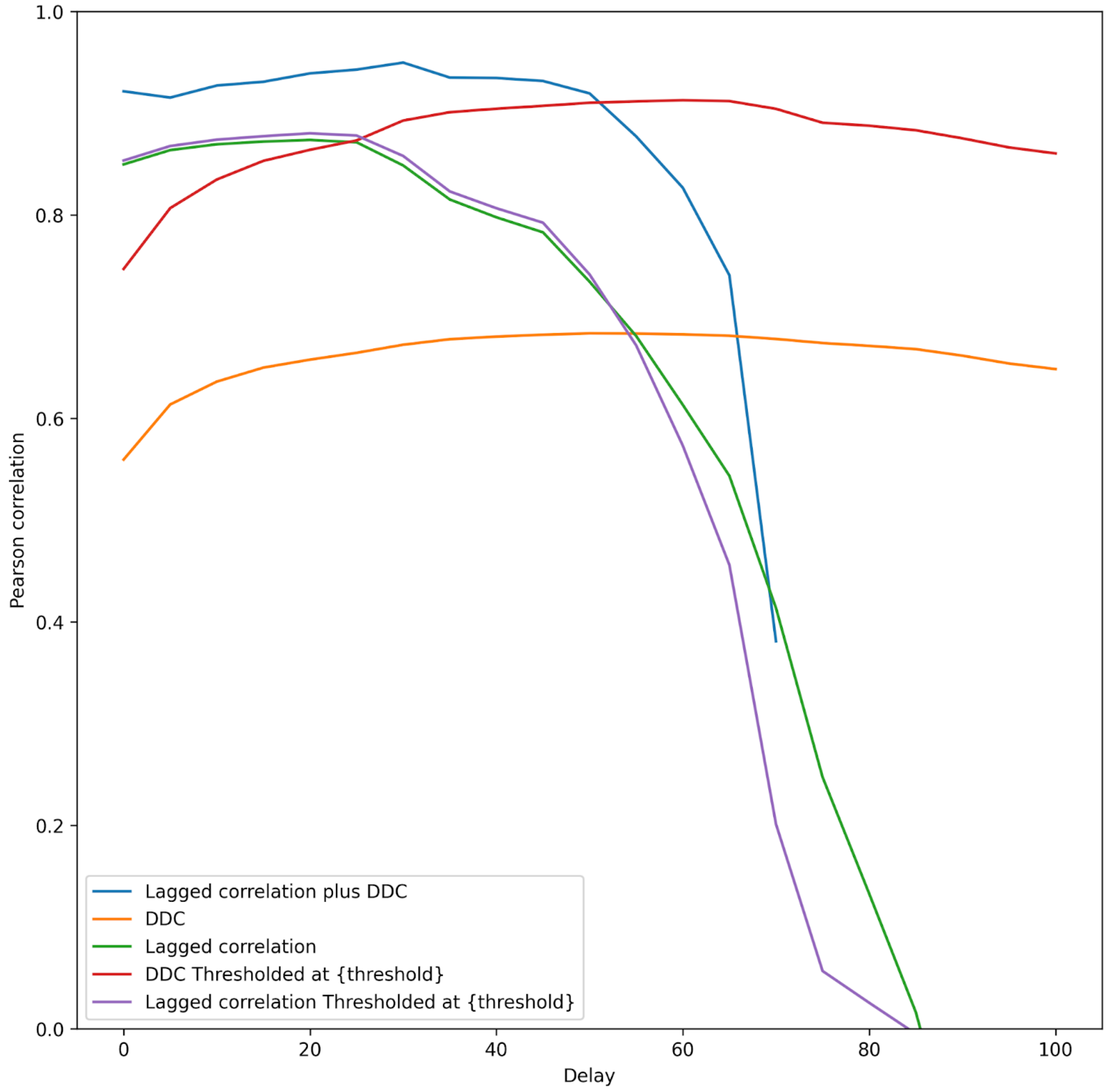


**Figure S12: Connectivity estimation performance measured by correlation of different methods on the Hopf model (N=10, p=0.1, a=0.25) with different fixed delays.** This plot shows the correlation of different methods estimating the connectivity matrix of a Hopf model with the delay parameter on the x-axis changing from 0 ms to 100 ms (fixed delays). Up to approximately a delay of 50 ms, Lagged Correlation in conjunction with DDC outperforms the other methods. Afterwards, the thresholded DDC algorithm is the only algorithm producing anything close to the actual GT connectivity matrix. N=10, p=0.1, a=0.25 (cf. SI Fig. 14).

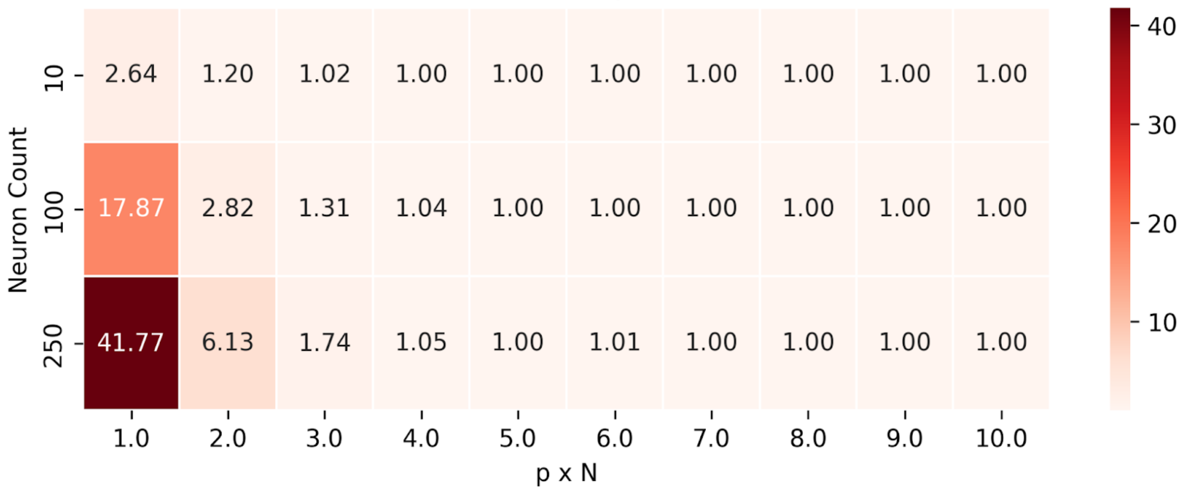


**Figure S13: Weakly connected components as a function of** $pN$**.**Weakly connected component analysis of the connectivity matrix in relation to neuron count and $pN$, the average number of inputs a neuron receives. The package networkx was used for the computation of weakly and also strongly connected components (https://networkx.org/documentation/stable/reference/algorithms/component.html).

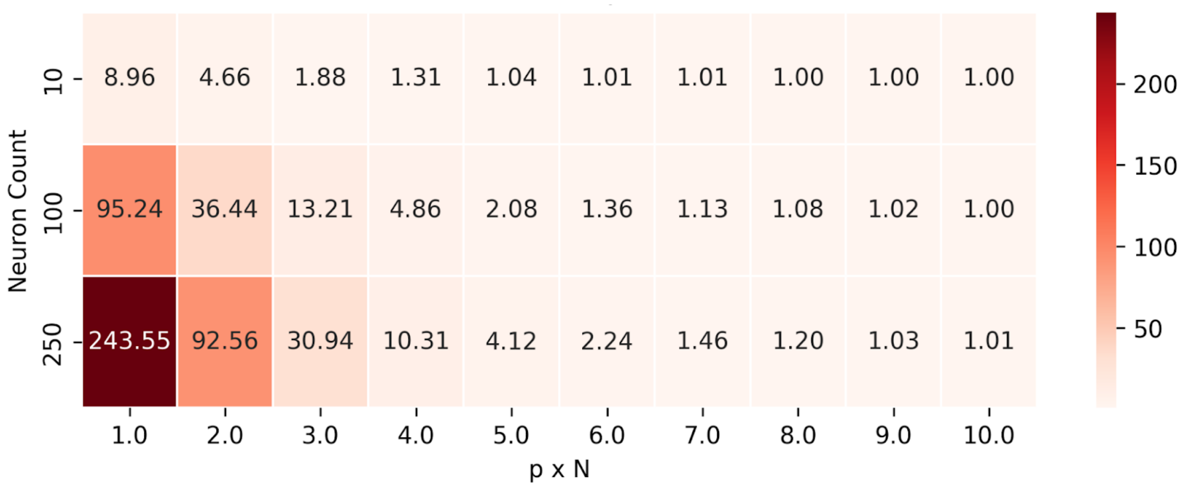


**Figure S14: Strongly connected components as a function of** $pN$**.** Strongly connected component analysis of the Connectivity matrix in relation to neuron count and $pN$, the average number of inputs a neuron receives.

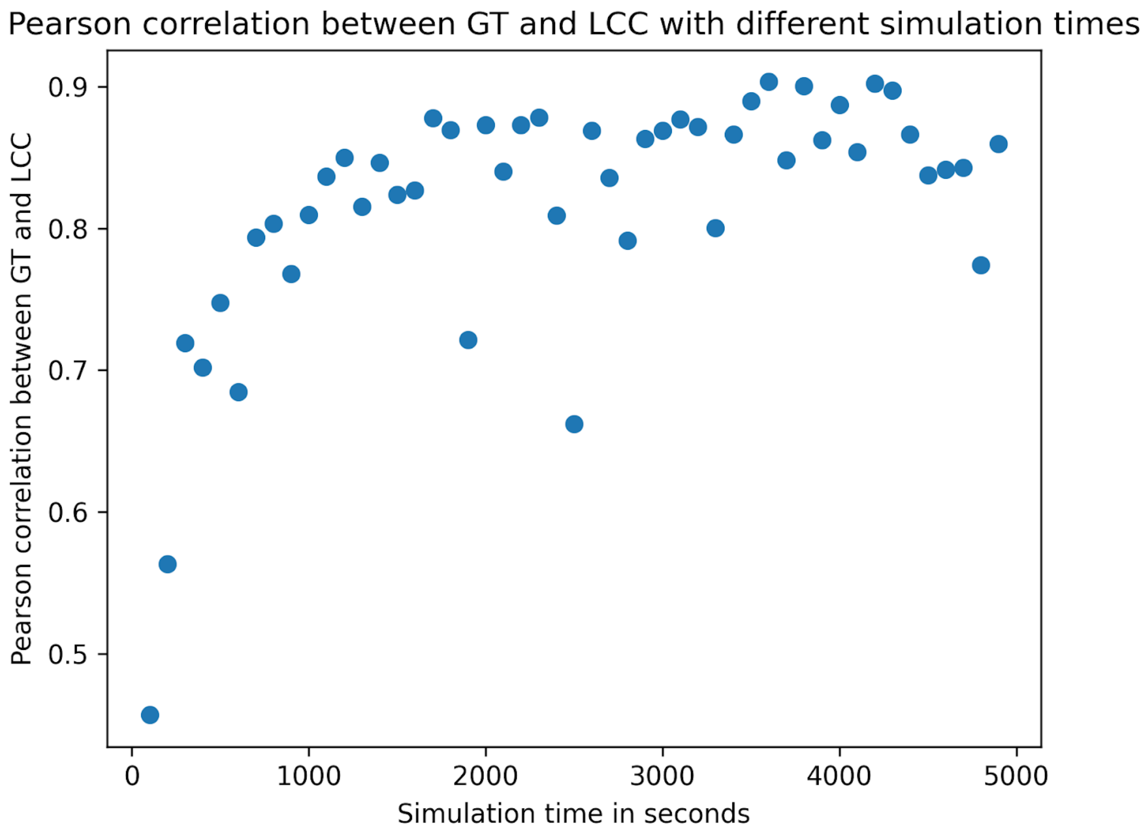


**Figure S15: Pearson correlation for different simulation times of the linear model.** Parameter values: $N = 100$ $p = 0.1$.

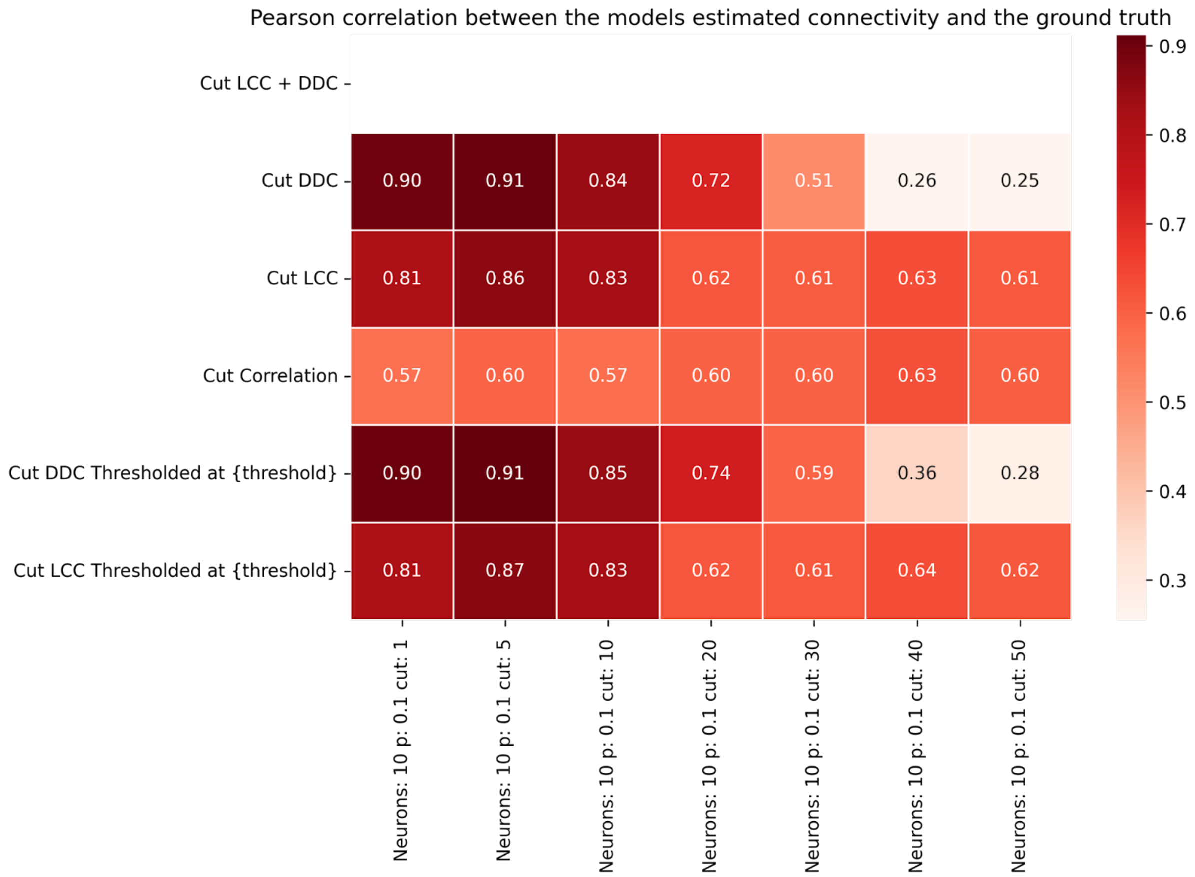


**Figure S16: Linear model (N=10, p=0.1): Connectivity estimation performance of cut methods in dependence on cut size for the linear model.** All parameters as in Fig. 5 of the main manuscript, except smaller M=100. Cut step size, which increases from left to right. All methods show lower performance with increasing cut step size.

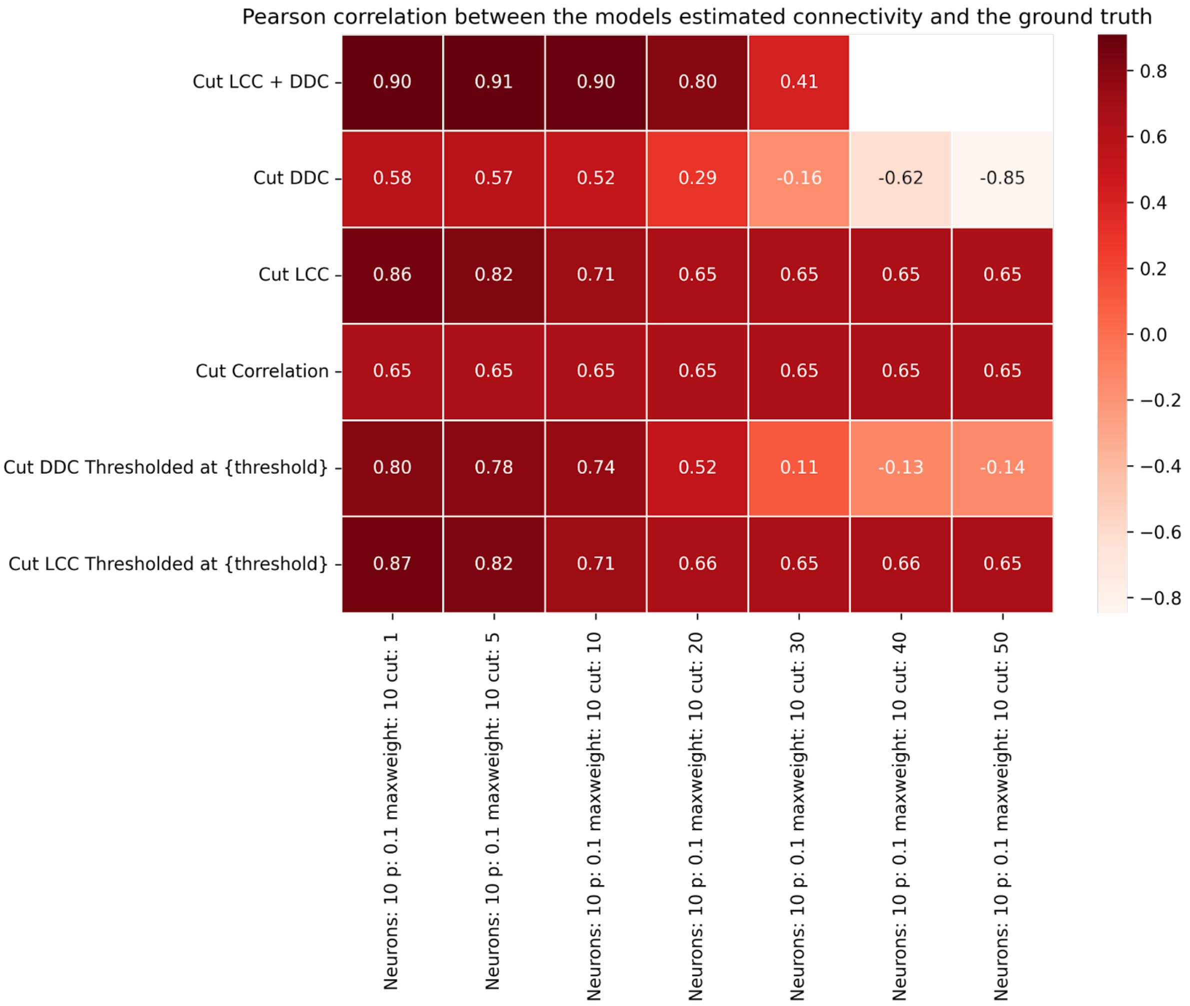


**Figure S17: Hopf model: Connectivity estimation performance of cut methods in dependence of cut size for small delays.** All parameters as in Fig. 2 in the main manuscript (N=10, p=0.1, a=0.25, delays in [0,10] ms). Cut step size increases from left to right. All methods show lower performance with increasing cut step size.

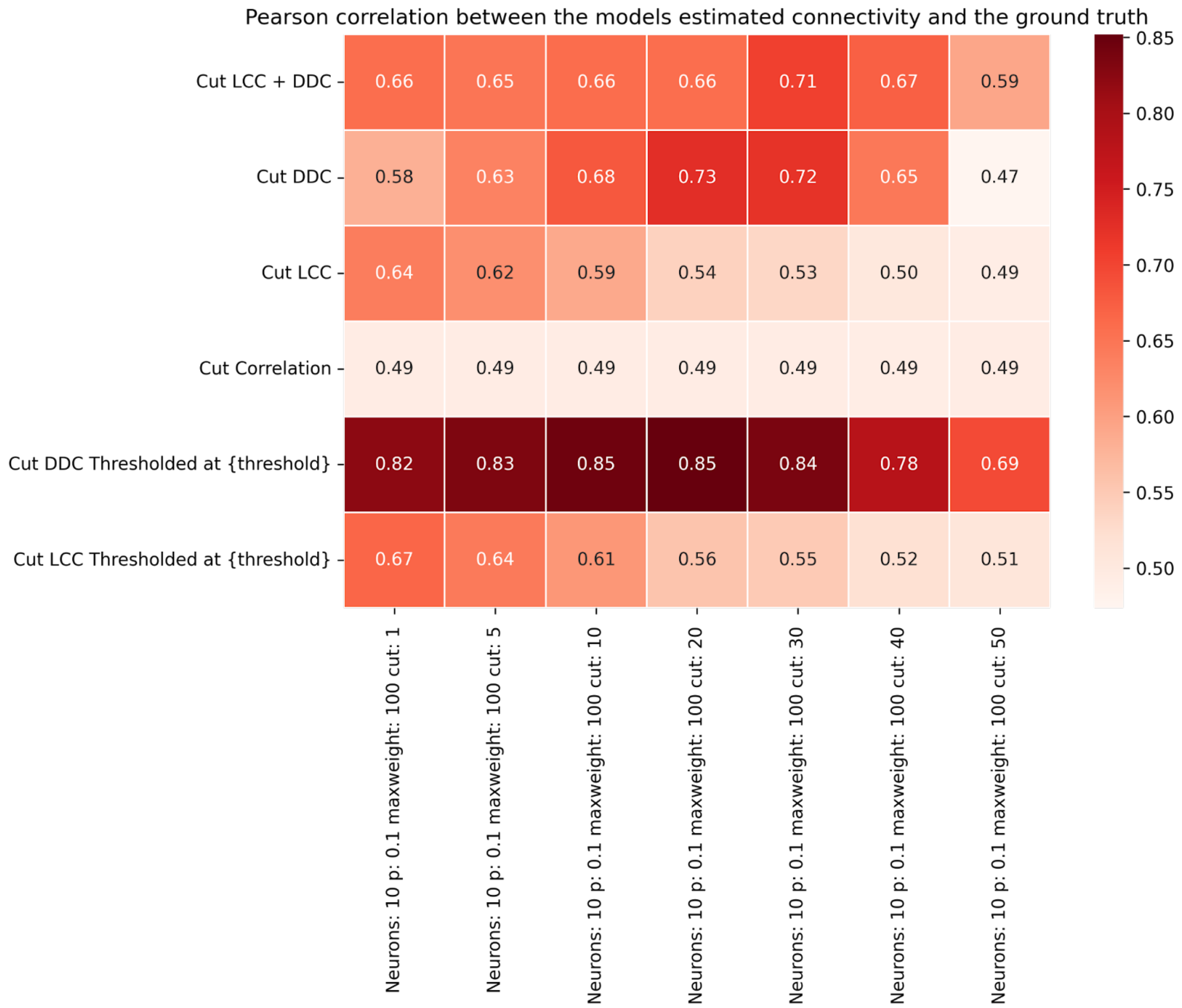


**Figure S18: Hopf model: Connectivity estimation performance of cut methods in dependence of cut size for larger delays.** All parameters as in Fig. 2 in the main manuscript, except for larger delays drawn uniformly between [0,100] ms. Cut step size, which increases from left to right. All methods, except Cut DDC thresholded and Cut DDC, show lower performance with increasing cut step size.

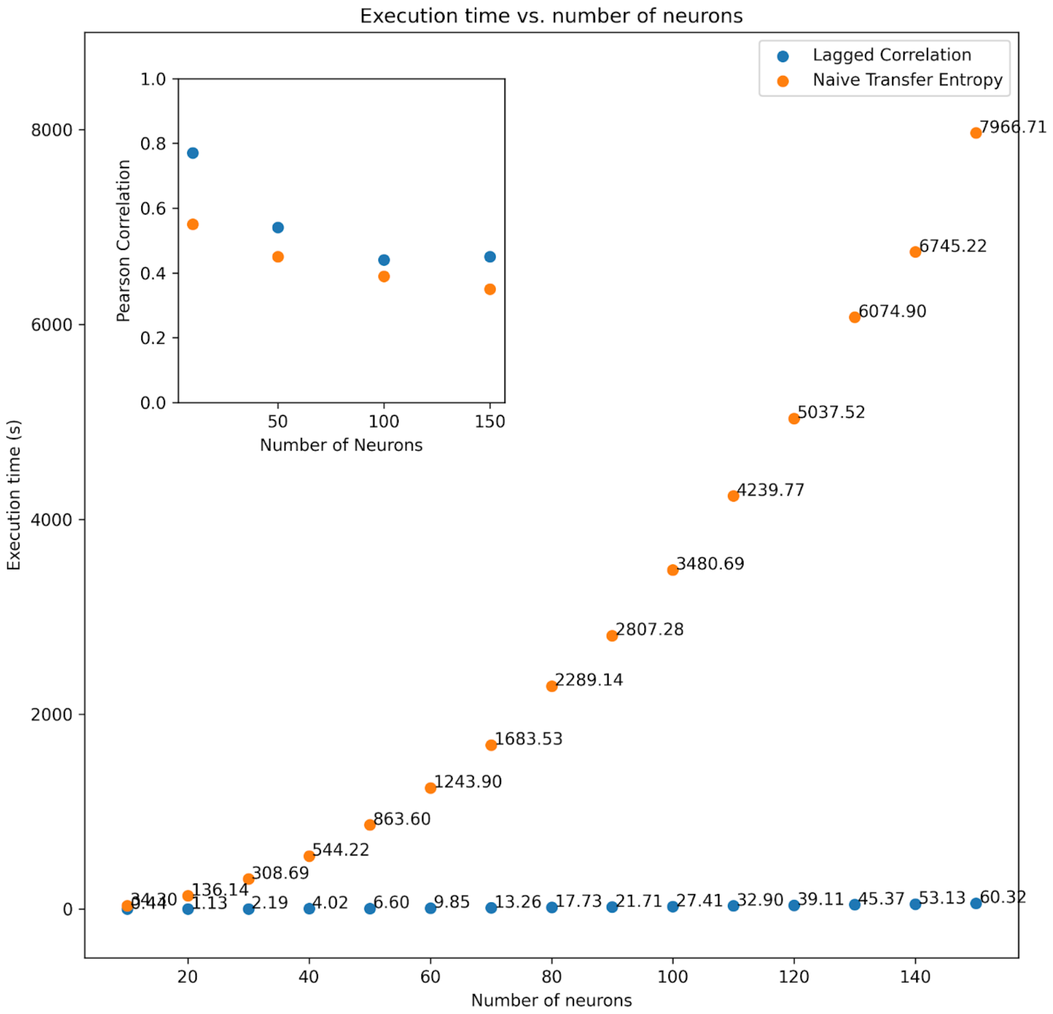


**Figure S19: Computation time and method performance as a function of number of neurons for LCC and NTE for the linear neuron model (Eq.8).** Inset plot shows the accuracy of both methods as quantified by Pearson correlation between GT at EStC matrices. LCC requires much lower computation time while showing a higher performance than NTE. Parameter value: $p = 0.1$.

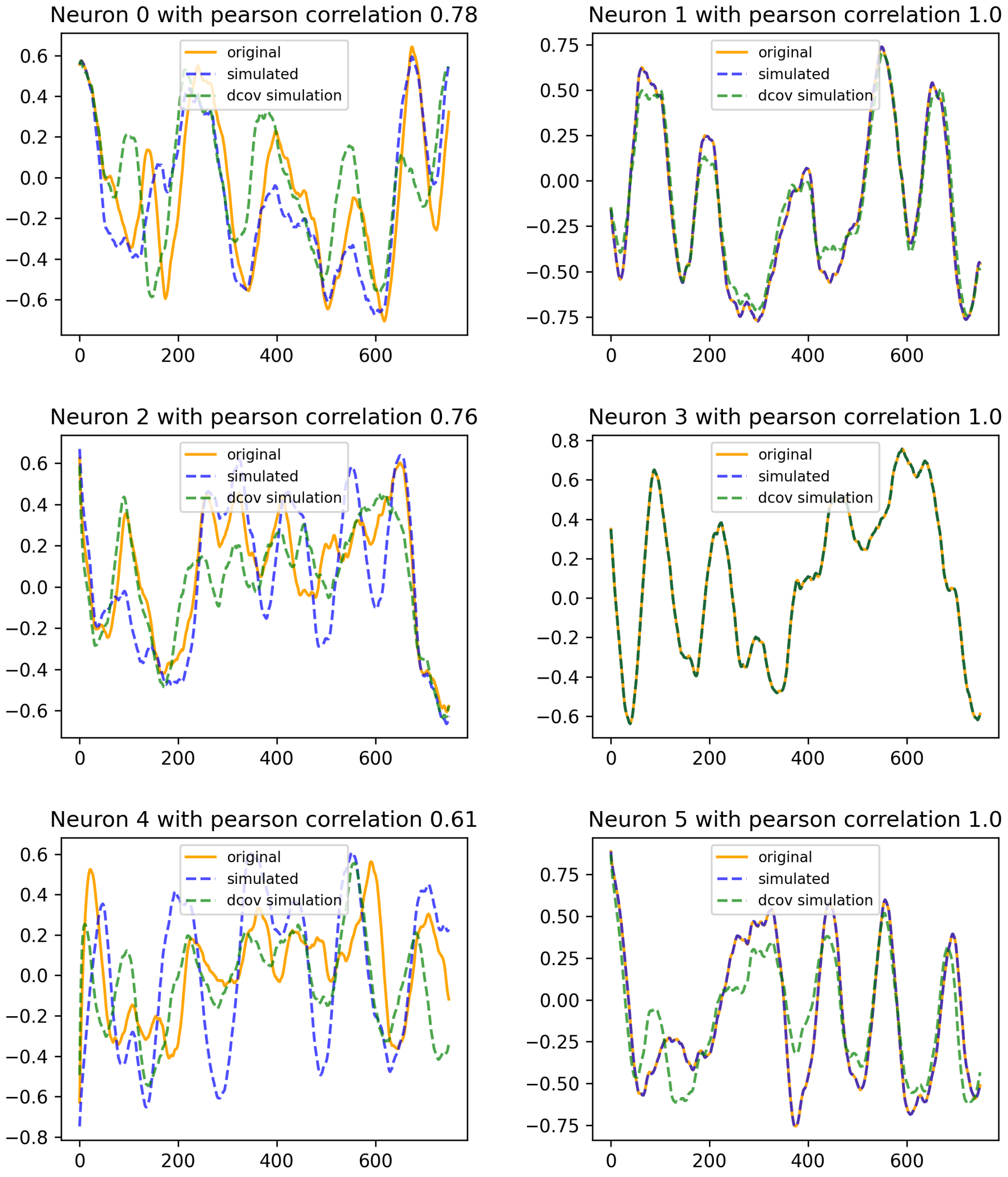
Neuron 0 with pearson correlation 0.78
original
simulated
dcov simulation
Neuron 1 with pearson correlation 1.0
original
simulated
dcov simulation
Neuron 2 with pearson correlation 0.76
original
simulated
dcov simulation
Neuron 3 with pearson correlation 1.0
original
simulated
dcov simulation
Neuron 4 with pearson correlation 0.61
original
simulated
dcov simulation
Neuron 5 with pearson correlation 1.0
original
simulated
dcov simulation

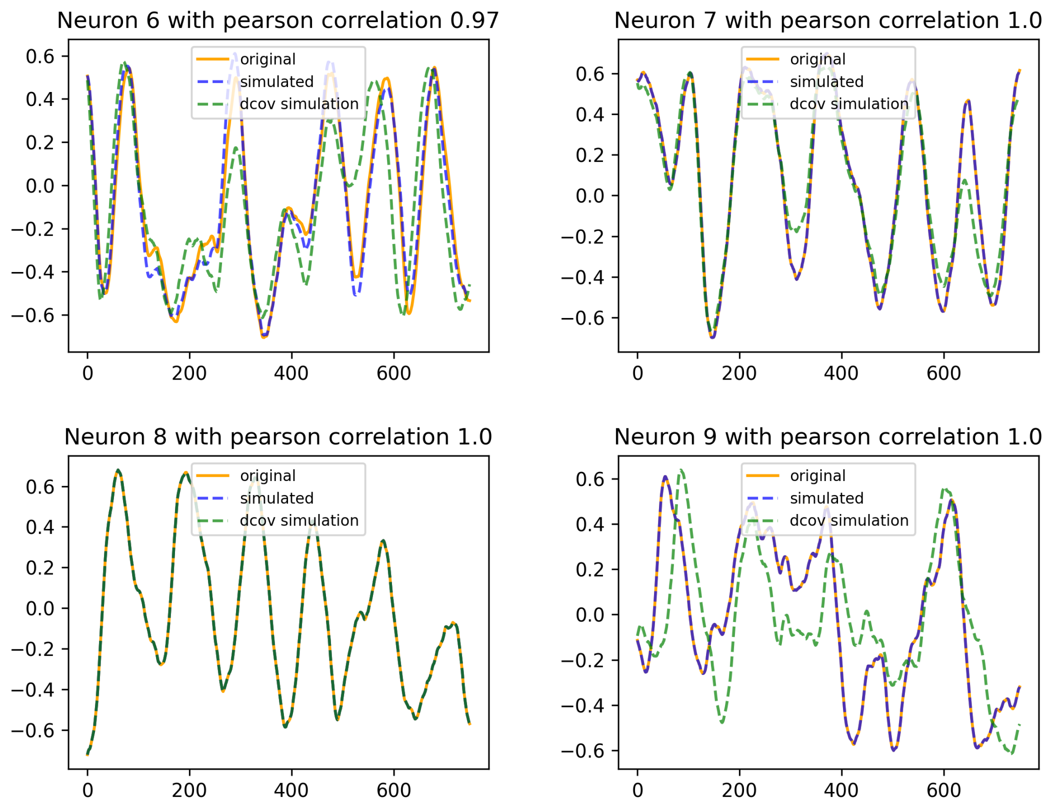


**Figure S20: Traces for all neurons in the Hopf model simulation and regeneration using the two estimation techniques.** The Pearson correlation is given for traces regenerated with LCC-estimated connectivity and GT traces.

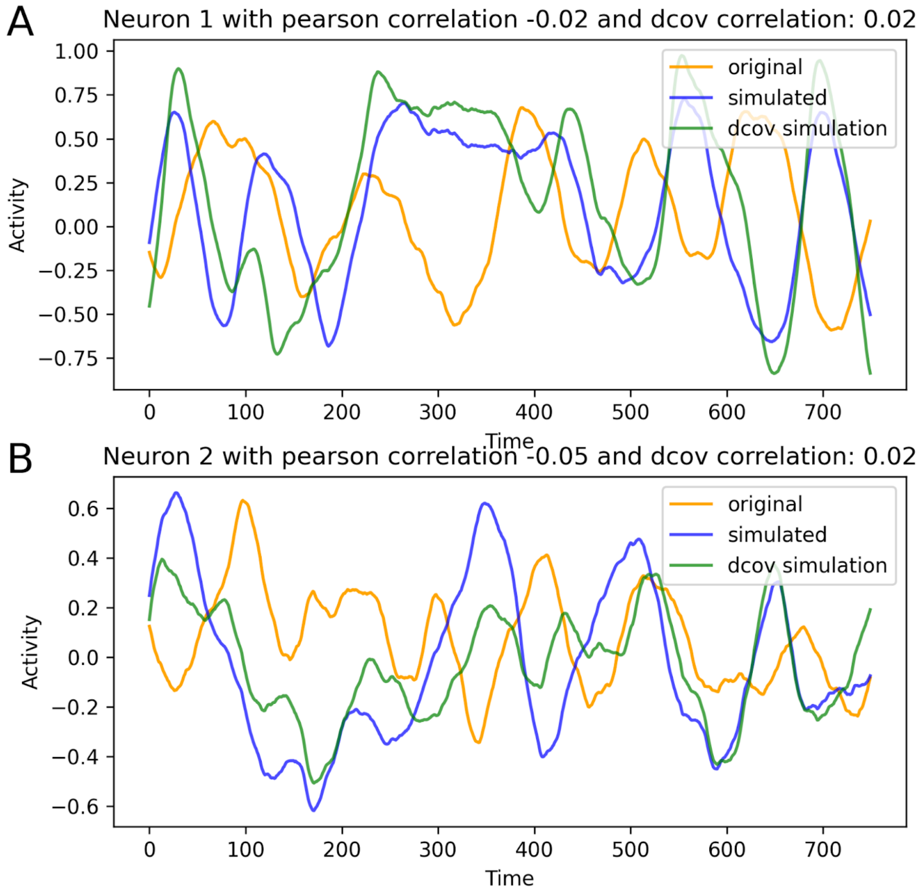


**Figure S21: Traces of the simulated Hopf Model when the seed for the LCC and DDC simulations were not the seed the original data was produced with.** Correlations between LCC-regenerated traces ('simulated') and DDC-regenerated traces ('dcov') are higher than for each method with the original traces.

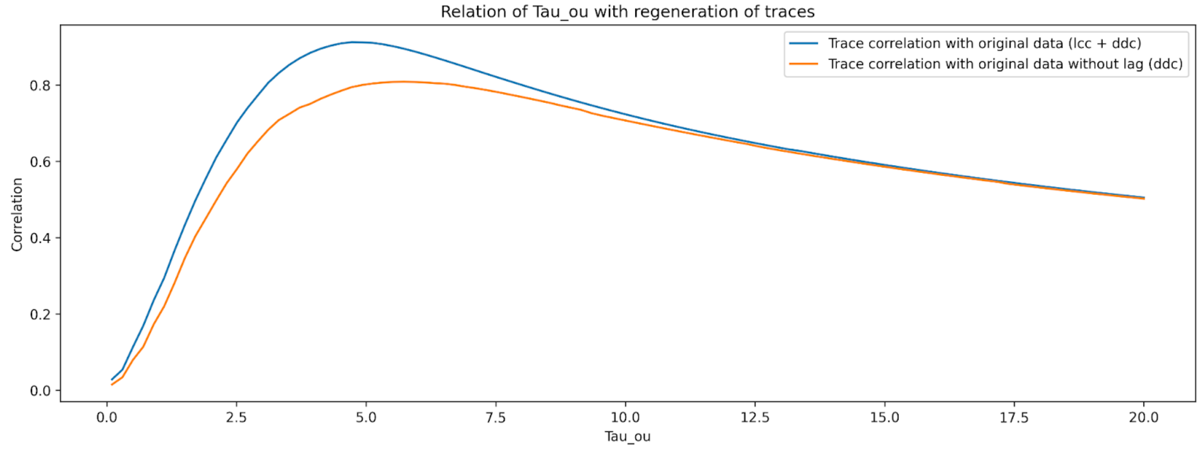


**Figure S22: Regeneration simulations: Influence of noise time constant tau_OU on the regeneration accuracy of the traces.** Shown is the mean trace-trace correlation (mean across neurons for original traces and regenerated traces using different estimation methods) for two estimation methods (LCC plus DDC and standard DDC). LCC plus DDC shows better performance than standard DDC. The mean trace correlation peaks at approximately tau_OU = 5 ms. Parameters as in Fig.8 in the manuscript.

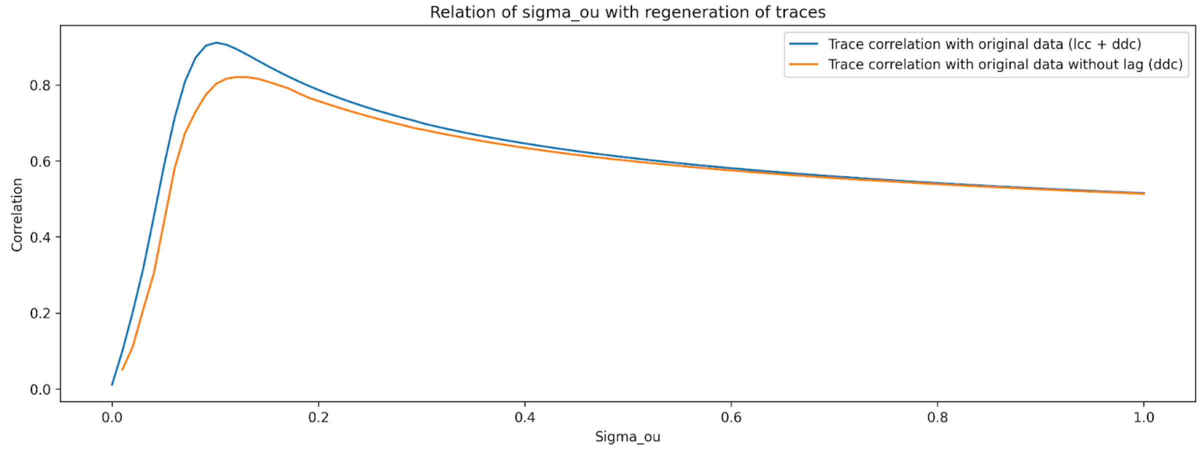


**Figure S23: Regeneration simulations: Influence of noise strength sigma_OU on the regeneration accuracy of the traces.** Figure layout as in SI Fig. 21. The mean trace correlation peaks at approximately sigma_OU = 0.1. Parameters as in Fig.8 in the manuscript.

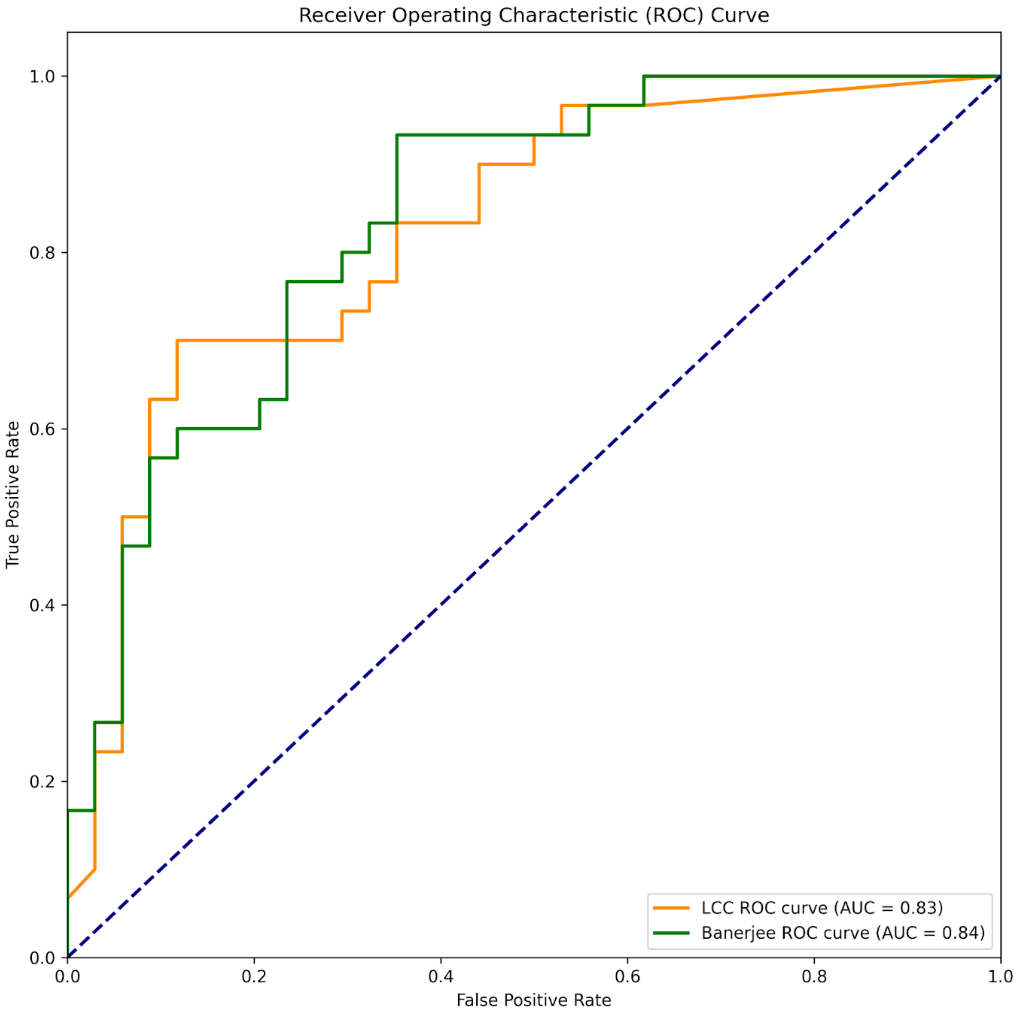


**Figure S24: ROC Curve comparing LCC and the reservoir computing method by Banerjeee et al. estimating connectivity of the *C. elegans* dataset.** Even though the reservoir computing method of Banerjee et al. has the larger AUC, the LCC algorithm outperforms it in the low false positive rates while still having a higher true positive rate.

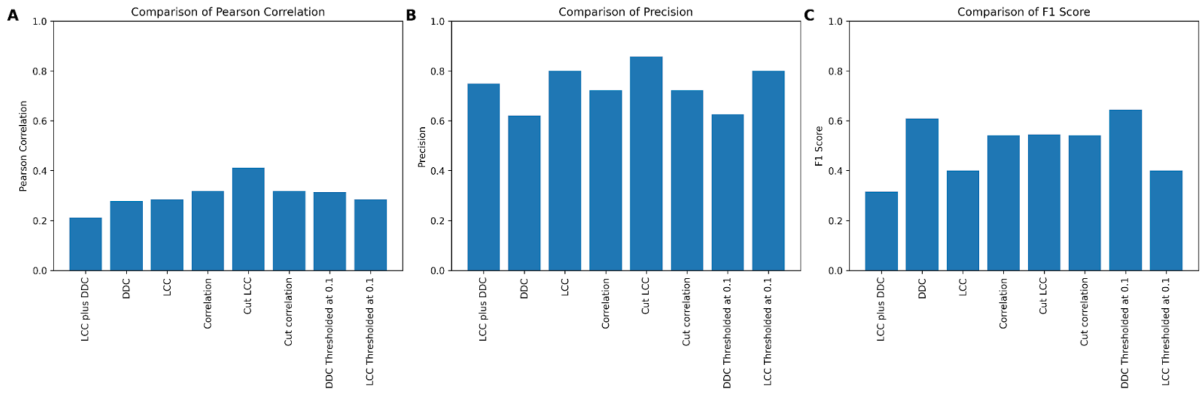


**Figure S25: Connectivity estimation performance of different methods on the *C. elegans* dataset.** The plot shows the correlation (panel A), the precision (panel B) and the F1 Score (panel C) of the estimated binary connectivity matrix for the *C. elegans* dataset. Cut Lagged Correlation outperforms the other methods in correlation and precision. For the F1 score, DDC and DDC thresholded perform better than the other methods. Overall, there is no pronounced difference between the methods.
